# GHOST: Recovering Historical Signal from Heterotachously-evolved Sequence Alignments

**DOI:** 10.1101/174789

**Authors:** Stephen M Crotty, Bui Quang Minh, Nigel G Bean, Barbara R Holland, Jonathan Tuke, Lars S Jermiin, Arndt von Haeseler

## Abstract

Molecular sequence data that have evolved under the influence of heterotachous evolutionary processes are known to mislead phylogenetic inference. We introduce the General Heterogeneous evolution On a Single Topology (GHOST) model of sequence evolution, implemented under a maximum-likelihood framework in the phylogenetic program IQ-TREE (http://www.iqtree.org). Simulations show that using the GHOST model, IQ-TREE can accurately recover the tree topology, branch lengths and substitution model parameters from heterotachously-evolved sequences. We develop a model selection algorithm based on simulation results, and investigate the performance of the GHOST model on empirical data by sampling phylogenomic alignments of varying lengths from a plastome alignment. We then carry out inference under the GHOST model on a phylogenomic dataset composed of 248 genes from 16 taxa, where we find the GHOST model concurs with the currently accepted view, placing turtles as a sister lineage of archosaurs, in contrast to results obtained using traditional variable rates-across-sites models. Finally, we apply the model to a dataset composed of a sodium channel gene of 11 fish taxa, finding that the GHOST model is able to infer a subtle component of the historical signal, linked to the previously established convergent evolution of the electric organ in two geographically distinct lineages of electric fish. We compare inference under the GHOST model to partitioning by codon position and show that, owing to the minimization of model constraints, the GHOST model is able to offer unique biological insights when applied to empirical data.

---

The success and reliability of model-based phylogenetic inference methods are limited by the adequacy of the models that are assumed to approximate the evolutionary process. Time-homogeneous models of sequence evolution have long been recognised as inadequate since the rate of evolution is known to vary across sites (Fitch and Margoliash, 1967; Holmquist et al., 1983) and across lineages (Lopez et al., 2002; Baele et al., 2006; Wu and Susko, 2011; Jayaswal et al., 2014). Many models have been proposed to compensate for rate heterogeneity across sites. The classical example is the discrete Γ model (Yang, 1994), which allows different classes of variable sites to have their rates drawn from a Γ distribution. More recently, Kalyaanamoorthy et al. (2017) relaxed the requirement for the rates of the classes to fit a Γ distribution, implementing a probability-distribution-free (PDF) rate model. However, these models still assume that the substitution rate for each site is constant across all lineages. This is too restrictive; biologically speaking it is not hard to accept that evolutionary processes can be both lineage and time dependent. In the context of a phylogenetic tree this manifests as lineage-specific shifts in evolutionary rate, coined heterotachy (Philippe and Lopez, 2001; Lopez et al., 2002), resulting in sequences that cannot be characterised as having evolved according to a single set of branch lengths and one substitution model.

The effect of heterotachy on phylogenetic inference was thrust into the spotlight by Kolaczkowski and Thornton (K&T) (2004). They used a simulation study to show that heterotachously-evolved sequences could mislead the popular inference methods of maximum-likelihood (ML) and Bayesian Markov Chain Monte-Carlo (BMCMC) to a greater extent than maximum parsimony (MP). Their findings were controversial and were widely challenged on the grounds that the simulations captured only a special case of heterotachy (Gadagkar and Kumar, 2005; Philippe et al., 2005; Spencer et al., 2005; Steel, 2005), and more general studies of heterotachy concluded that ML performed at least as well as, and in most cases better than, MP (Gadagkar and Kumar, 2005; Spencer et al., 2005). Valid as these criticisms may have been, the key issue that the K&T study brought to light stood firm - heterotachy was a primary source of model misspecification and the models and methods of the time were ill-equipped to deal with it.

The main impediment to the development of models that can accommodate heterotachously-evolved sequences has been the computational expense. Models that account for heterogeneity of rates of change across sites can be integrated relatively cheaply, but modeling heterotachy is not so simple. One approach has been covarion (COV) models (Fitch and Markowitz, 1970). Tuffley and Steel (1998) described a model in which sites could switch between variable and invariable states in different lineages. All variable sites in the model shared a common substitution model and rate. This model was gradually extended (Galtier, 2001; Huelsenbeck, 2002), eventually reaching its most complex form in which sites can switch along lineages between a number of different rates as well as an invariable state (Wang et al., 2007).

Another approach has been to use partition models (Lanfear et al., 2012), which require the data to be partitioned *a priori*. The analysis then proceeds by inferring seperate branch length and model parameters for each partition. Sequence data are commonly partitioned based on genes and/or codon position. However, the inherent assumption of this approach is that heterotachy only occurs between partitions, not within each partition. This may not be a valid assumption, so the requirement to partition the data in advance of the analysis is a possible source of model misspecification.

An alternative approach has been to use mixture models, in which the likelihood of the data at each site in the alignment is calculated as a weighted sum across multiple classes (see Pagel and Meade (2005) for a detailed description of phylogenetic mixture models). The most common approaches can be referred to as mixed substitution rate (MSR) models (Lartillot and Philippe, 2004; Pagel and Meade, 2004; Foster, 2004), whereby each class has its own substitution rate matrix; and mixed branch length (MBL) models (Kolaczkowski and Thornton, 2004; Meade and Pagel, 2008), whereby each class has its own set of branch lengths on the tree. Hybrid versions of these models have also been proposed, such as the HAL-HAS model of Jayaswal et al. (2014). Zhou et al. (2007) compared a covarion model to an MBL model, finding the covarion model to be more efficient at handling heterotachy. They did however conclude that both methods warranted further exploration, going on to propose the covarion mixture model (CM) (Zhou et al., 2010), which incorporates covarion parameters that vary across sites. As a consequence of their parameter rich nature, these models have all been implemented only within a Bayesian framework. Wu and Susko (2009) proposed a general framework for heterotachy, encompassing both MSR and MBL models as special cases. Another example is the CAT models of Lartillot and Philippe (2004), which have been widely used (Whelan and Halanych (2017) and references therein). Whelan and Halanych (2017) carried out extensive simulation and empirical studies comparing the performance of the CAT models to partition models. They concluded that despite their additional complexity and associated increase in runtime, the CAT models generally perform no better than partition models. They also lamented that when new mixture models are introduced in the literature their performance is not always assessed against the current popular methods for phylogenetic analysis, such as partition models.

As a consequence of their varied nature, mixture models require many parameters and the associated computational expense has thus far impeded their implementation in a ML framework. The issue of computational expense is an ever diminishing one; as computing power increases and algorithmic architecture improves, the opportunity to employ more and more complex models of sequence evolution does also. We introduce the General Heterogeneous evolution On a Single Topology (GHOST) model for ML inference. The GHOST model combines features of both MSR and MBL models. It consists of a number of classes, all evolving on the same tree topology. For each class, the branch lengths, nucleotide or amino-acid frequencies, substitution rates and class weight are all parameters to be inferred. The motivation behind this modelling philosophy is the desire to minimise assumptions that might lead to model misspecification. Although the cost of this philosophy, in terms of model complexity and the associated risk of over-parameterisation, is not to be ignored, by refraining from placing strict constraints on the inference we allow the opportunity to recover new, and perhaps surprising, historical signals from the data. We provide an easy-to-use, ML implementation of the GHOST model in the phylogenetic program IQ-TREE (Nguyen et al., 2015) (http://www.iqtree.org), the first mixture model of comparable flexibility to be made available in a ML framework.

## Methods and Materials

### Model Description

The GHOST model consists of a user-specified number of classes, *m*, and one inferred tree topology, *T*, common to all classes. All other parameters are inferred separately for each class. For the *j*^*th*^ class, we define λ_***j***_ as the set of branch lengths on *T*; ***R***_***j***_, the relative substitution rate parameters; ***F***_***j***_, the set of nucleotide or amino-acid frequencies; and *w*_*j*_, the class weight (*w*_*j*_*>* 0*, ∑w*_*j*_ = 1). Given a multiple sequence alignment (MSA), *A*, we define *L*_*ij*_ as the likelihood of the data observed at the *i*^*th*^ site in *A* under the *j*^*th*^ class of the GHOST model. *L*_*ij*_ is computed using Felsenstein’s (1981) pruning algorithm. The likelihood of the *i*^*th*^ site, *L*_*i*_, is then given by the weighted sum of the *L*_*ij*_ over all *j*:

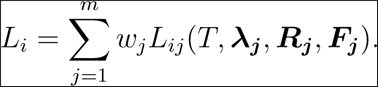

Therefore, if *A* contains *N* sites (length of the alignment), the full log-likelihood, *l*, is given by:

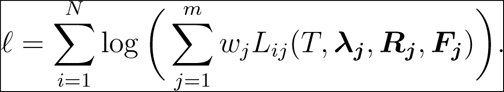

We make use of the existing parameter optimisation algorithms within IQ-TREE, extending them, where necessary, to incorporate parameter estimation across the *m* classes.

### Model Parameter Estimation for a Fixed Tree, T

Given a fixed tree topology, *T*, let **Θ** = *{w*_1_*,…, w*_*m*_, ***λ*_1_***,…*, ***λ***_***m***_, ***R*_1_***,…*, ***R***_***m***_, ***F***_1_,…, ***F***_***m***_*}* denote the GHOST model parameters (*i.e.*, class weights, branch lengths, relative substitution rates, and nucleotide or amino-acid frequencies) for each of the *m* classes. To estimate all parameters for a tree, *T*, we employ the expectation-maximization (EM) algorithm (Dempster et al., 1977; Wang et al., 2008). We initialize **Θ** with all 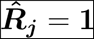 in each class, uniform nucleotide or amino-acid frequencies 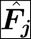 (i.e., the Jukes-Cantor model), and ŵ_*j*_ and 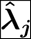 obtained by parsimonious branch lengths rescaled according to the rate parameters of a discrete, PDF rate model (Kalyaanamoorthy et al., 2017) with *m* categories. This becomes the current estimate 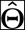. The EM algorithm iteratively performs an expectation (E) step and a maximization (M) step to update the current estimate until an optimum in likelihood is reached.

### Derivation of Expectation-Maximization algorithm

The premise underlying the GHOST model is that each site evolved according to just one of the *m* classes, however we do not have any information about which sites belong to which class. We define **c** = *{c*_1_*, c*_2_*,…, c*_*N*_*}* as a vector that maps the *N* sites to one of the *m* classes. The EM algorithm works by formulating an expression for the expected value of our objective function and then maximizing that expectation. In the context of GHOST, we can restate the likelihood equation as follows:

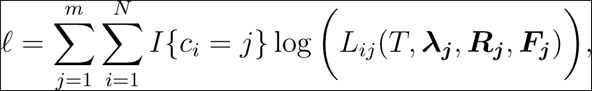

where *I{c*_*i*_ = *j}* is an indicator function that is equal to 1 when the class of the *i*^*th*^ site is equal to *j*, and 0 otherwise. Taking the expectation of this expression yields:

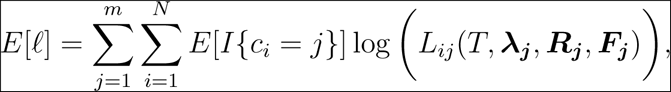

#### E-step

In the context of the GHOST mixture model, the goal of the E-step is to evaluate the quantity *E*[*I{c*_*i*_ = *j}*] for a fixed set of tree and model parameters. An intuitive interpretation of the expected value of this indicator function, is that it is simply the probability that a given site *i* belongs to a given class *j*. For simplicity, we define this quantity as 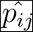 and evaluate it using a simple application of Bayes Theorem. Given the current parameter estimates, we can calculate 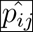 as follows:

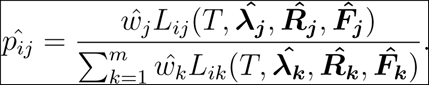

#### M-step

The goal of the M-step is then to update the parameter estimates to maximize the expected likelihood, fixing the *p*_*ij*_ that were calculated during the E-step. For each class *j*, we maximize the expectation of the log-likelihood function:

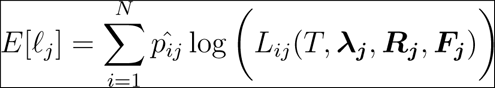

to obtain the next 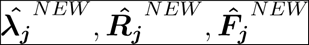. Within IQ-TREE, 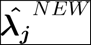is obtained via Newton-Raphson optimization, while 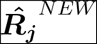and 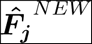are estimated by the Broyden-Fletcher-Goldfarb-Shanno (BFGS) algorithm (Fletcher, 2013). Finally, the weights are updated by:

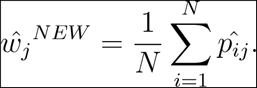

That is, the new weight for class *j* is the mean posterior probability of each site belonging to class *j*. This completes the proposal of the new estimate 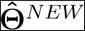 If 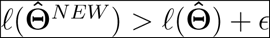 (where *ϵ* is a user-defined tolerance, *ϵ* = 0.01 by default), then 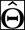is replaced by 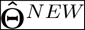 and the E and M steps are repeated. Otherwise, the EM algorithm finishes.

An auxiliary benefit of the ML implementation of the GHOST model in IQ-TREE is that once the EM-algorithm has converged, we can soft-classify sites to classes, according to their probability of belonging to a particular class. This classification can be used to identify sites in the alignment that belong with high probability to a particular class of interest.

### Tree search

The tree search algorithm in IQ-TREE (Nguyen et al., 2015) is based on the construction of a candidate tree set. Trees from the candidate tree set are rearranged by Nearest Neighbour Interchange (NNI) to explore the tree space. This algorithm was tested extensively during the ML implementation of the GHOST model and two significant changes to the heuristic were required:

1. In the original implementation of IQ-TREE, after each NNI is performed, a single branch length optimization step for the five branches adjacent to the NNI is carried out. We found that this amount of branch length optimization was insufficient for the GHOST model. Instead, IQ-TREE now performs *m* branch length optimization steps after each NNI (where *m* is the number of classes in the GHOST model).
2. Prior to the ML implementation of the GHOST model, IQ-TREE only fully optimised the model parameters of the best tree in the candidate tree set. During the ML implementation of the GHOST model we found that this technique proved to provide too much of an advantage to the current best tree. The algorithm was modified such that when the GHOST model is used, all trees in the set of candidate trees are fully optimized.

### Software

The GHOST model has been implemented in IQ-TREE (Nguyen et al., 2015) (http://www.iqtree.org). A list of commands for use of the GHOST model in IQ-TREE can be found in the supplementary material (available on the Dryad data repository, doi: TBA).

### Testing of the GHOST Model in IQ-TREE

We tested the efficacy of the ML implementation of the GHOST model in IQ-TREE by carrying out three separate simulation studies. The first study was a replication of the simulations carried out by Kolaczkowski and Thornton (2004), focusing on IQ-TREE’s ability to recover the correct tree topology from heterotachously-evolved data on quartet trees. We found that using the GHOST model, IQ-TREE was able to reliably recover the simulation parameters in all cases. The methods, results (Supplementary Figs. S1 and S2) and discussion of these simulations can be found in the supplementary material. The second study used 12-taxon trees and focused on IQ-TREE’s ability to recover branch length and substitution model parameters from heterotachously-evolved data. The third study used 32-taxon trees and focused on the establishment of a sound model selection procedure, specifically to determine the correct number of classes from simulated alignments. Finally, we investigated the effect of using the incorrect number of classes on topological accuracy.

#### 12-taxon simulations

The replication of the Kolaczkowski and Thornton simulations focused on recovering tree topology only. However, the GHOST model is parameter rich and naturally the implementation process must assess the ability of IQ-TREE to accurately recover branch lengths and model parameters under the GHOST model. We constructed independent sets of parameters for two classes on a randomly generated 12-taxon tree using the GTR model of evolution. For each class the branch lengths were drawn randomly from an exponential distribution with a mean of 0.1. We then used *Seq-Gen* (Rambaut and Grassly, 1997) to simulate MSAs. When specifying a GTR rate matrix in Seq-Gen, the G*↔*T substitution rate is fixed at 1 and all other substitution rates are expressed relatively. Within each class, the five relative substitution rates were drawn randomly from a uniform distribution between 0.5 and 5. The four base frequencies for each class were assigned a minimum of 0.1, with the remainder allocated proportionally by scaling a normalised set of four observations from a uniform (0, 1) distribution. From these two classes MSAs were constructed by varying the weight of each class. The weight of Class 1, *w*_1_, was varied from 0.2 to 0.8 in increments of 0.05 and at each increment 20 separate MSAs were simulated. Each MSA was constructed by concatenating two independently simulated sets of sequences, the first of length 10, 000 *× w*_1_ simulated using the Class 1 parameters, and the second of length 10, 000 *×* (1 *− w*_1_) simulated using the Class 2 parameters. We used IQ-TREE to infer parameters from each MSA under a GHOST model with two GTR classes (GTR+FO*H2). We also inferred parameters from each MSA under a partitioned GTR model, where the branch length parameters were unlinked (*i*.*e*., estimated separately for each partition). We also repeated the procedure with a range of shorter sequence lengths: 100, 500, 1000, and 5000 nucleotides.

The accuracy of inferred base frequency and relative rate parameters for the 12-taxon simulations was measured by calculating the mean absolute difference between the inferred and true parameters. The accuracy of branch length estimates was assessed using the branch score metric, BS (Kuhner and Felsenstein, 1994). In order to assess the accuracy of branch length recovery we needed to establish a frame of reference to gauge whether the results obtained are suitably close to the truth or not. To do this we made use of the estimates under the branch-unlinked partition model as a baseline. The fundamental difference between the partition model and the GHOST model is that the partition model has *a priori* knowledge of which sites in the alignment belong to which class. This means that in effect (and excluding the possibility of inferring the incorrect topology) the results of the partition model are identical to those that would be obtained by fitting GTR models to the Class 1 and Class 2 sequences independently. Naturally we cannot expect that the GHOST model can perform better than this, so we can consider the accuracy of the partition model as a benchmark.

### Model selection

#### 32-taxon simulations

In order for the GHOST model to be used on empirical sequence alignments we must have some method of model selection, in particular selecting the appropriate number of classes. Information criterion methods such as Akaike’s Information Criterion (AIC) (Akaike, 1974) or Bayesian Information Criterion (BIC) (Schwarz et al., 1978) are commonly used in phylogenetics for this purpose, so we carried out simulations to establish whether these criteria could accurately predict the correct number of classes that generated the alignment. We also investigated the influence of the number of classes inferred on topological accuracy. We generated 300 heterotachous sequence alignments for each of *m* = 2, 3 and 4 classes. Each alignment was 10,000 bp long, contained 32 taxa and used the GTR model of sequence evolution for each class. The weight of each class, *w*_*i*_, was held fixed at 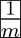. For each alignment, the model parameters for each of the *m* classes were generated as in the 12-taxon simulations. For each alignment, a ‘base set’ of branch lengths, ***λ***, was generated randomly from an exponential distribution with a mean of 0.1. The branch length parameters for the *m* classes were then generated as follows:

1. For the *i*^*th*^ class, a vector of random variables (of same length as ***λ***), **s**_**i**_, was drawn from a uniform distribution on (0, 1).
2. For the *i*^*th*^ class, a class scaling factor, *α*_*i*_, was drawn from a uniform distribution on (0, 1).
3. Finally, an overall scaling factor, *β*, was calculated to ensure that the weighted total tree length (TTL) of the *m* classes was equivalent to the TTL of the ‘base set’:

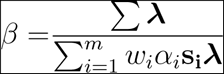
4. The branch length vectors for the *i*^*th*^ class were then given by:

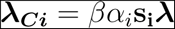

For the *i*^*th*^ class, we used *Seq-Gen* to simulate a sequence alignment of length 10,000*×w*_*i*_ bp. These were concatenated together to form the heterotachous alignment. This procedure was repeated to generate 300 heterotachous sequence alignments for each of *m ∈ {*2, 3, 4*}*.

For each of the 900 simualted alignments we used IQ-TREE to fit GHOST models with 1, 2, 3*,…*, 8 classes. For each alignment, we used AIC and BIC (where we used sequence length as the proxy for *n* in the BIC formula) to determine the number of classes that provided the best fit between tree, model and data. In order to select between models when AIC and BIC do not agree, we examined the inferred parameters when too many classes were used to see if we could recognise characteristic signs of overfitting. We also investigated the influence of the inferred number of classes on the topological accuracy, as measured by the Robinson-Foulds (RF) distance (Robinson and Foulds, 1981). Finally, we investigated the computation time required for IQ-TREE to arrive at ML estimates under the GHOST model, as a function of the number of classes in the model (Supplementary Figure S3).

#### Plastome alignments

In order to investigate the variability in the number of classes recommended by AIC and BIC on empirical alignments, we created seperate empirical alignments by subsampling from a plastome alignment, taken from Yan et al. (2017), which consisted of 66 genes for 26 species. We discarded all genes shorter than 1000 bp, leaving a total of 20 genes. From these 20 genes, we randomly sampled 20 groups of 1, 3, 5, 10 and 15 genes to create a total of 100 separate alignments. We then fitted GHOST models with increasing number of classes to each alignment to determine the number of classes that provided the best fit according to both AIC and BIC.

### Placement of Turtles Among Archosaurs

One can think of the linked version of the GHOST model in terms of the discrete Γ model, with the removal of some constraints. The linked GHOST model does not require the classes to be of equal weight, nor does it impose that the branch lengths between classes are correlated. The PDF rate model can be thought of as an intermediate step between the discrete Γ and the linked GHOST models. To demonstrate the effect of relaxing these constraints we applied 4-class discrete Γ, PDF rate and linked GHOST models to a phylogenomic alignment consisting of 248 genes (187,026 bp) for 16 taxa. The alignment was taken from Chiari et al. (2012), in which they concluded that turtles were a sister group to birds and crocodiles, as opposed to crocodiles only.

### Convergent Evolution of the Na_v_1.4a Gene Among Teleosts

We applied the GHOST model to a sequence alignment (2178 bp) taken from the coding region of a sodium channel gene, Na_*v*_1.4a, for 11 teleost species.

Model selection is the first challenge when using the GHOST model on an empirical alignment. We tested a wide variety of substitution models, as shown in Supplementary Figure S4. Starting with the two-class GHOST model, we used IQ-TREE to optimise the likelihood of the data under each substitution model. Subsequently, we repeated the process with up to a maximum of six classes. For each run we used the unlinked version of the GHOST model, so that each class had its own set of branch lengths, base frequencies and substitution model parameters inferred. We then applied our information theory-based model selection criteria to determine the substitution model and number of classes that provided the best fit. For the best GHOST model, we also tested the linked versions to evaluate whether inferring model parameters individually for each class was necessary. Finally, we found the best PDF rate model (Kalyaanamoorthy et al., 2017) and compared that to the best GHOST model based on AIC.

In order to compare the GHOST model to alternative current phylogenetic methods, we also used IQ-TREE to fit a branch-unlinked partition model. The electric fish alignment was split into three partitions, based on codon structure. We then used PartitionFinder (Lanfear et al., 2012) to evaluate the best substitution models to use on each partition. Finally, IQ-TREE was used to fit the best branch-unlinked partition model to the alignment, using the models of sequence evolution suggested by PartitionFinder.

## Results & Discussion

### 12-taxon simulations

We simulated heterotachously-evolved MSAs of varying lengths (100, 500, 1,000, 5,000 and 10,000 bp) on a random 12-taxon tree topology, with two classes evolvingaccording to the GTR model of evolution. Figure 1 shows the performance of the GHOST model in recovering the various tree and model parameters for Class 1 of the 10,000 bp simulated alignments. The analagous plots for all sequence lengths and both classes can be found in Supplementary Figures S5-S12. The results of the 12-taxon simulations show that under the GTR+FO*H2 model IQ-TREE recovered the base frequencies, relative rate parameters and weights to a high degree of accuracy for both classes. With respect to the branch score (BS) (Fig. 1c and Supplementary Figs. S9 and S10), we see that the GHOST model again performs well. The mean BS for the GHOST model approaches that obtained by the partition model as class weight (and therefore share of sequence length in the mixture) increases, despite the partition model enjoying the advantage of having full knowledge of which sites were simulated under which class. A BS of zero would imply that the true simulation parameters were inferred for every simulated alignment. Thus, the magnitude of the BS for the partition model can be thought of as a measure of the stochastic simulation error. The difference between the BS for the GHOST and partition models can then be considered the error attributable to losing the knowledge of the partitioning scheme. This error appears negligible in comparison to the simulation error. In Figure 1c, when *w*_1_ *>* 0.5, the overlap of the error bars (which represent *±*2 standard errors of the mean) suggests that the trees inferred by the GHOST model are not significantly different from those inferred by the partition model. This is a promising result, as in empirical data any partitioning of the MSA is based on assumptions, and therefore introduces a potential source of model misspecification. The GHOST model can be applied without any such assumptions.

**Figure 1:**
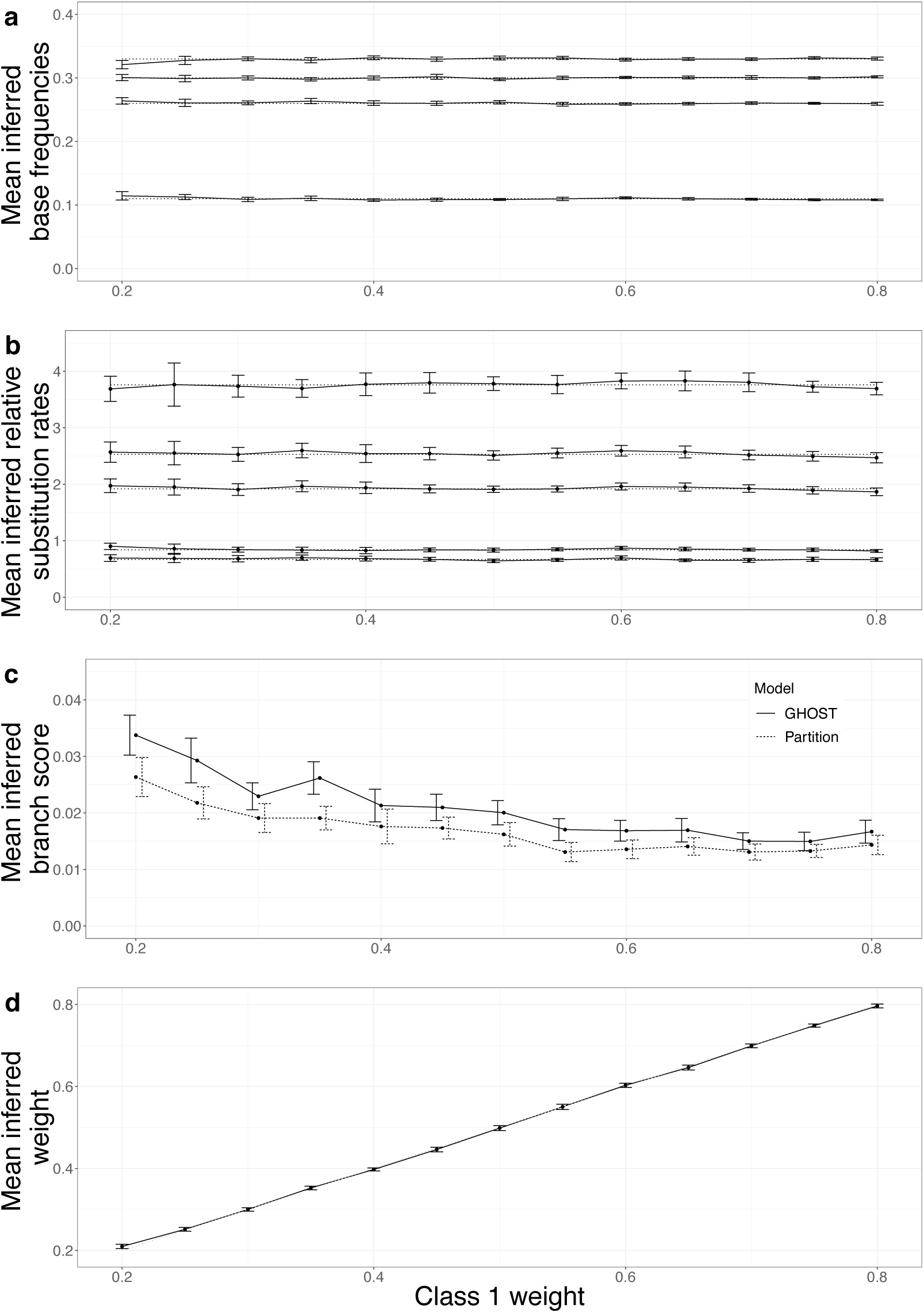
12-taxon simulations, 10,000 bp alignments - Class 1 inferred parameters vs Class 1 weight. The data points indicate the mean value of the inferred parameter or statistic, the error bars represent *±*2 standard errors of the mean. Dotted lines represent the true parameter value used for data simulation. (a) Base frequencies (b) Relative substitution rates (c) Branch score (BS) for both the GHOST and partition models (d) Inferred Class 1 weight.

To demonstrate the ability of the GHOST model to provide meaningful information about which sites might belong to which class, we performed a soft classification on one of the MSAs generated for the 12-taxon simulations. For simplicity we have chosen an MSA where Class 1 and Class 2 are of equal weight. Supplementary Figure S13 indicates that the probability of a site belonging to Class 1 is generally higher for those sites that were simulated under the Class 1 parameters. However, given the stochastic element of the simulations, there are some sites simulated under the Class 2 parameters that are classified as having a higher probability of evolving under Class 1, and *vice versa*. For this reason, we never attempt to ‘hard classify’ the sites, that is, allocating specific sites to a particular class with absolute certainty. Rather, we ‘soft classify’ the sites, that is, we consider that a site belongs to all classes, according to its probability distribution of evolving under each class.

#### The effect of sequence length

An important consideration when employing parameter rich models is the amount of information in the alignment. Estimating many parameters from an insufficient amount of information will result in unreliable parameter estimates. Supplementary Figure S14 shows that the GHOST model and the partition model recover the correct tree topology at similar rates. For simulated alignments of length 100 bp, tree inference was poor for both GHOST (30.8% inferred trees correct) and the partition model (33.5%). This failing is quickly remedied by increasing sequence length, with topological accuracy for both models greater than 90% for 500 bp alignments. When looking at parameter inference we see a similar story. Supplementary Figures S5 to S12 show the progressive improvement in the accuracy of inferred parameters as sequence length is increased. For sequence lengths of 100 bp the parameter estimates are completely unreliable, as is the inferred topology. This is not surprising given the dearth of information on which to base the inference. As sequence length increases so does the strength of the phylogenetic signal from each class. At 500 and 1,000 bp, the estimates are reasonably close to the true values but still exhibit a moderate level of variance. For 5,000 and 10,000 bp the parameter estimates are very close to the true values and with little variance. These 12-taxon, 2-class simulations have a total of 59 free parameters to be estimated. Based on theseresutls it would seem prudent when applying the GHOST model to empirical datasets to ensure a minimum of 10*k* sites in the alignment, where *k* is the number of free parameters under the proposed model.

### Model Selection

#### 32-taxon simulations

The primary purpose of the 32-taxon simulations was to establish a sound model selection technique to allow the GHOST model to be applied to empirical alignments with confidence. Information theory methods such as AIC and BIC are typically used by phylogeneticists to choose amongst models. How these two methods perform on complex mixture models such as GHOST is unclear. Zhou et al. (2007) found that when applied to models with high numbers of parameters, AIC tended to overfit the data (inclusion of parameters is penalised too lightly) whereas BIC tended to underfit the data (inclusion of parameters is penalised too heavily). Dziak et al. (2018) counsel that while information criteria are useful guides, they do have their limitations, and so nuance and judgment remain important elements in the model selection process.

For each of the 900 simulated alignments (300 for each *m ∈ {*2, 3, 4*}*, 10,000 bp long), we used AIC and BIC to determine the optimal number of classes for IQ-TREE to infer under the GHOST model. The results are summarised in Table 1. AIC selects the correct number of classes in 95% of cases for *m* = 2, always erring on the side of overfitting. As *m* increases, the accuracy of AIC rises to more than 99% for *m* = 4. BIC selects the correct number of classes 100% of the time for *m* = 2, but the accuracy of BIC decreases as *m* increases, dropping to 90% for *m* = 4. Conversely to AIC and in line with expectations based on the literature, BIC always erred on the side of underfitting.

**Table 1:**
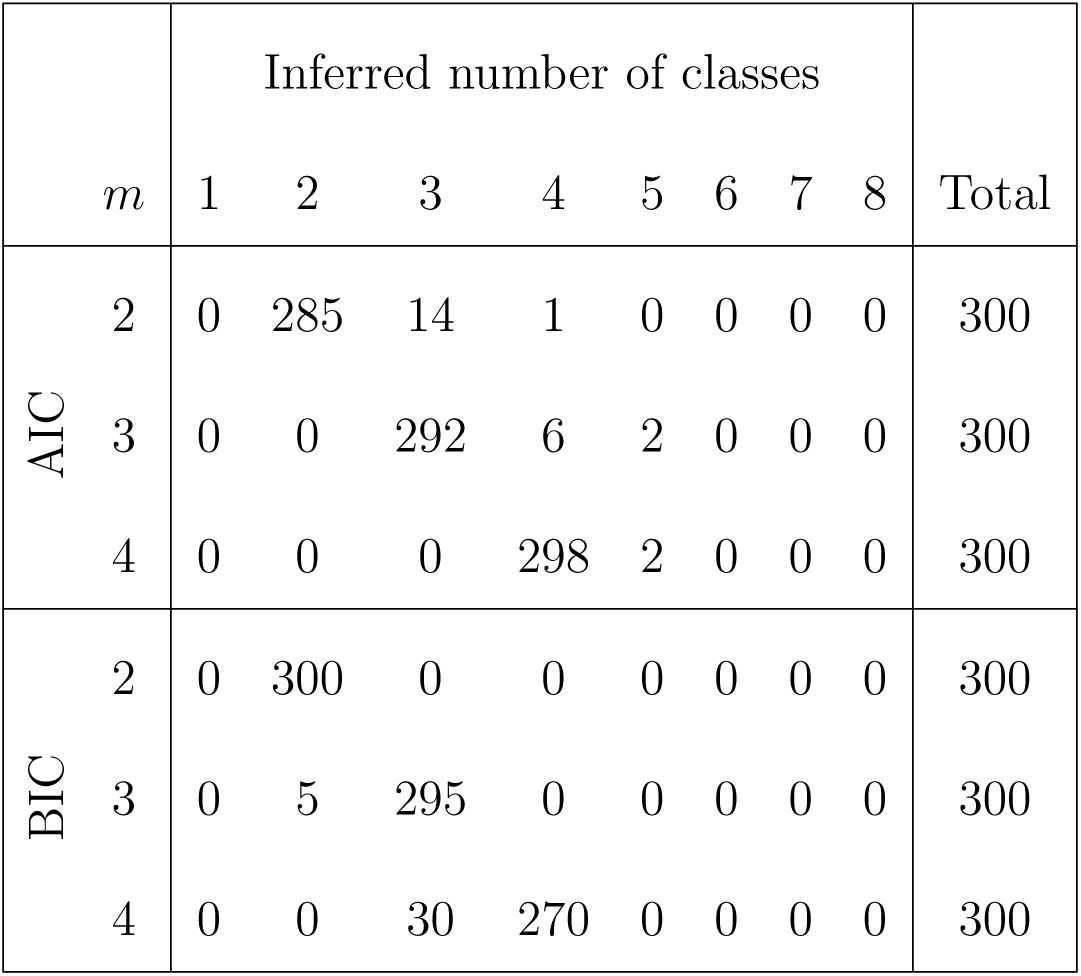
32-taxon simulations, model selection using AIC or BIC. For each of the 900 simulated alignments (300 for each *m* {*∈*2, 3, 4}, where *m* is the true number of simulated classes), we used AIC and BIC to determine the optimal number of classes to infer under the GHOST model.

#### Plastome alignments

The results of the 32-taxon simulations discussed above indicate that BIC and AIC agree on the number of classes in the vast majority of cases, so there is little ambiguity in the model selection process. However, this may not be the case in empirical alignments. We subsampled genes from a phylogenomic alignment to create 100 different alignments, 20 each of single-gene, 3-gene, 5-gene, 10-gene and 15-gene alignments. Supplementary Figure S15 shows the level of variability between the number of classes recommended by BIC and AIC. It is apparent that the broad agreement between BIC and AIC when applied to simulated alignments is not mirrored in empirical data. One reason for this might be that when applied to the simulated alignments, the true model is available as one of the candidate models and so both criteria tend to select this model or something quite close to it. This is obviously not the case for empirical data, and so this may explain why we see considerably more variation in the results between the criteria. Regardless, it does highlight that when applying the GHOST model to empirical alignments, choosing the number of classes requires a more nuanced approach to be developed.

#### Choosing the number of classes

Model selection can be thought of as a trade-off between bias (the chosen model has too few parameters to adequately represent the underlying evolutionary processes) and variance (the model has too many parameters to provide stable parameter estimates) (Burnham and Anderson, 2003; Posada and Buckley, 2004). Given that the primary motivation behind the development of the GHOST model was the minimization of model misspecification, we should prefer modest overfitting to modest underfitting. A model that has too many classes has the advantage that the true model is nested within it, and therefore the true parameters remain recoverable, albeit with some undesirable redundancy. Conversely, a model with too few classes must merge at a minimum two classes into one, and therefore the true parameters are not recoverable. Thus, we can respectively consider the BIC and AIC-based optimal number of classes as a lower and upper bound on the number of classes in the best-fit GHOST model. The challenge is to find a way to sensibly choose the optimal number of classes between these bounds.

Intuitively, there does not seem to be any way to predict the effect of underfitting (fitting less classes than was used to generate the data) on the inferred parameters. However, the same is not true of overfitting. If we fit too many classes then we may expect one of two things to happen:

1. We will recover the true branch lengths, model parameters and weights for the correct number of classes, with any remaining classes having weight very close to zero.
2. We may have two or more inferred classes in which the inferred branch lengths and model parameters are very similar to each other, with the sum of their weights being approximately equal to the weight of a single true class.

To investigate whether the above propositions hold, we examined the results of the 32-taxon, 4-class simulations. We compared results when the correct, 4-class GHOST model was used vs. the overfit, 5-class GHOST model. Each of the 300 alignments were simulated from 4 classes of equal weight, so we decided to look at the weight of the inferred classes as a proxy for successful recovery of the true classes. It would seem an unlikely coincidence if we inferred four classes of approximately equal weight without them closely resembling the true classes in terms of branch lengths and model parameters. Supplementary Figure S16 shows the variability in inferred class weights when the correct number of classes (GTR+FO*H4) is used. With the exception of a few outliers, all the inferred classes have a weight close to the simulated value (0.25). If we add an extra class such that the model is overfitted (GTR+FO*H5) as in Supplementary Figure S17, we see that the three largest inferred classes have weight close to the true value of 0.25, with the remaining weight split between the two smallest classes. The rightmost box in Figure S17 shows the sum of the weights for the two smallest classes, which is again consistently close to 0.25. These observations are consistent with the propositions outlined above, in which three of the four simulated classes are inferred with reasonable accuracy, while the fourth simulated class is split into two inferred classes. To further explore this hypothesis, we checked if the branch lengths inferred in the smallest class were more strongly correlated with the branch lengths of the second smallest inferred class than those of the three largest classes. We expect this effect to be stronger as the weight of the smallest inferred class increases. Supplementary Figure S18 shows two matrices, displaying the correlation between the branch lengths inferred by the five classes. The classes are ordered by weight, C1 referring to the largest class and C5 referring to the smallest.

Supplementary Figure S18 (a) shows the correlation for alignments in which the inferred weight of C5 was less than 0.05, and we see that for these cases (138 of 300 alignments) the branch lengths of C5 and C4 have a correlation of 0.32; whereas shows the correlation among those alignments for which it was greater than 0.05, and we see that for these cases (162 of 300 alignments) the correlation is much higher at 0.79. Based on this evidence, we can highlight two characterisitic signs of overfitting, that users can check for when choosing the number of classes to fit to empirical data:

1. One or more of the inferred classes has a negligible weight in comparison with the other classes.
2. The trees of two or more of the inferred classes show strong similarities in terms of branch lengths.

It is possible then to recommend the following approach when selecting the number of classes to fit to empirical alignments:

1. Calculate the maximum number of classes that is reasonable to use for a given alignment. The criterion that should be used is that the number of free parameters in the model must be no more than 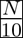, where N is the length of the alignment. Call this number *U.*
2. Without exceeding *U*, find the optimal number of classes as judged by both BIC (call this number *L*) and AIC (if this is less than *U*, then update *U* accordingly). We now have a lower (*L*) and upper (*U*) bound on the number of classes that should be considered as potentially providing the best fit.
3. Examine the class weights and trees inferred by the GHOST model with *U* classes. If none of the class weights are negligible and the trees are all reasonably distinct then accept *U* as the optimal number of classes for the GHOST model.
4. If signs of overfitting are present, examine the class weights and trees for the GHOST model with *U* – 1 classes. If no overfitting is present then accept this number of classes as optimal.
5. Repeat Step 4, continually removing classes until no signs of overfitting are present, or until *L* is reached.

The forthcoming discussion of the convergent evolution of the Na_*v*_1.4a is an example of an empirical alignment in which AIC appears to give a reasonable number of classes, with no signs of overfitting present. A counter example is provided Crotty et al. (2018), where AIC is found to overfit the data whereas BIC offers a more reasonable fit.

#### Impact of model misspecification

While we consider the model selection procedure outlined above to be reasonable, it must be remembered that it is not deterministic and it was developed based on the performance of the GHOST model on simulated alignments. We must therefore recognise the potential for over/underfitting to occur in practice with empirical alignments, and assess the potential impact of such errors. To do so we used the 32-taxon simulations to investigate the effect of choosing the wrong number of classes on IQ-TREE’s ability to infer the correct topology under the GHOST model. We calculated the RF distance between the trees used for simulation and those inferred by IQ-TREE. Figure 2 displays the mean RF distance as a function of the number of classes in the fitted model, expressed relative to *m*, the true number of classes used to simualte the alignments. As we should expect, for all values of *m* the mean RF distance is minimized when *m* classes are inferred. However, the mean RF distance increases much faster in the presence of underfitting than it does in the presence of overfitting. This finding supports the use of the top-down approach when choosing the number of classes as described above, as any errors will tend to be on the side of overfitting rather than underfitting. Detailed summaries of the distribution of RF distances are given in Supplementary Tables S1, S2 and S3 for *m* = 2, 3 and 4, respectively.

**Figure 2:**
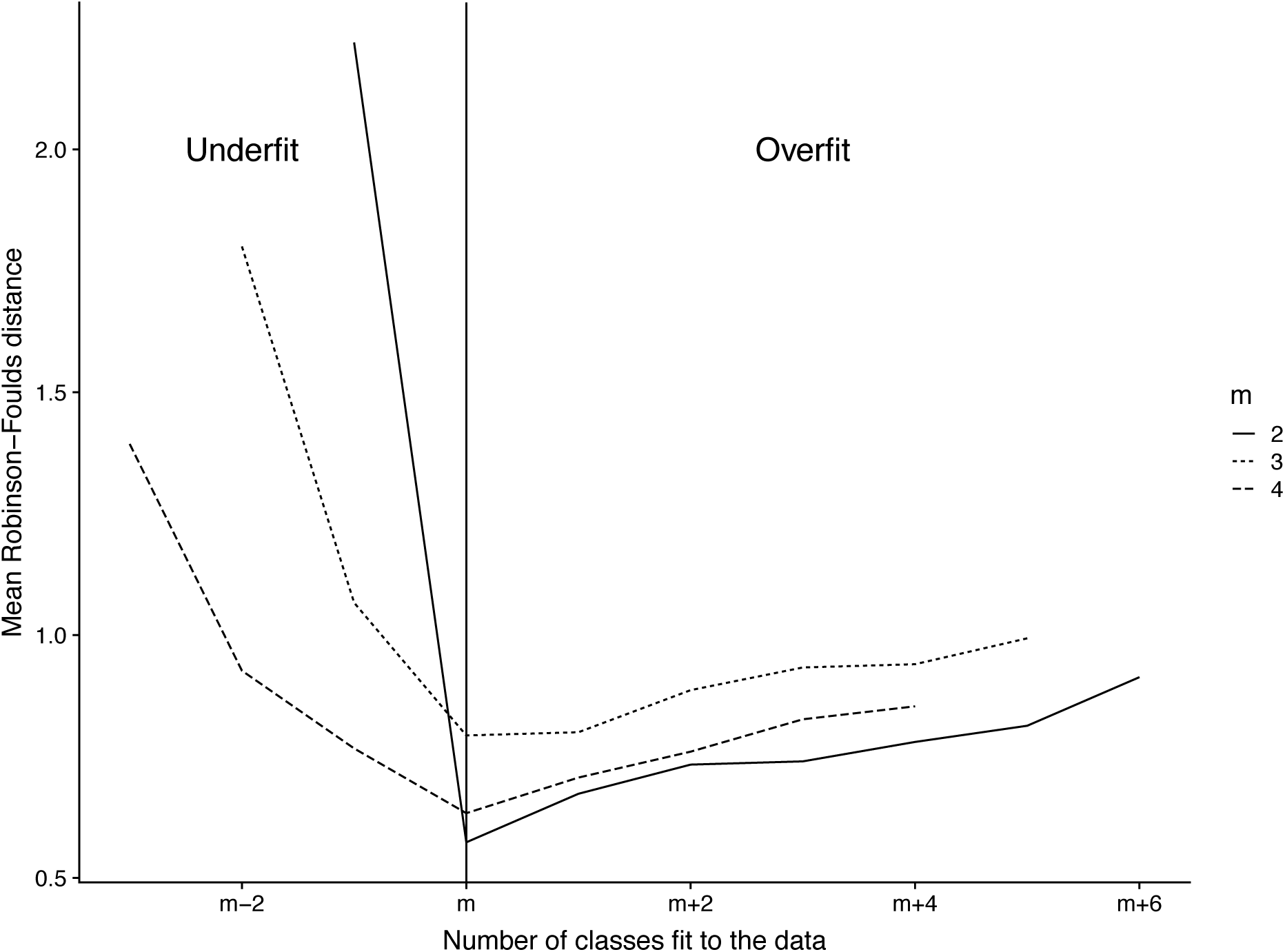
32-taxon simulations, effect of under/overfitting on topological accuracy, for the 900 simulated alignments (300 each for *m* ∈ {2, 3, 4}). The y-axis displays the mean RF distance between the inferred trees and the trees used to simulate the alignment. The x-axis shows the number of classes used for the inference, expressed relative to *m*, the true number of classes used to simulate the alignments.

### Placement of Turtles Among Archosaurs

The placement of turtles in the phylogenetic tree of amniotes has been controversial, due in part to their morphological peculiarities. It is currently accepted that turtles are a sister lineage to archosaurs (birds and crocodiles), as opposed to crocodiles alone. Chiari et al. (2012) assembled and analyzed a 248-gene, 187,026 nucleotide alignment of 16 taxa, concluding that the tendency to place turtles as sister to crocodiles was a phylogenetic artefact caused by saturation at codon position 3 sites. They found the preferred grouping of turtles as sister to archosaurs was returned when the alignment was partitioned by codon position or when only codon position 1 and 2 sites were included. Among the models that returned the non-preferred topology was the GTR+G, with four rate categories. To examine the influence on this result of the restrictions imposed by the discrete Γ model, we tested the discrete Γ, the PDF rate model and the GHOST model on the same alignment. In order to ensure a fair comparison all models used four classes (as in Chiari et al. (2012) and the linked version of the GHOST model was used. Supplementary Table S4 indicates that the GHOST model proved superior in terms of both AIC and BIC. The resulting tree topologies can be found in Figure 3, showing that the discrete Γ and PDF rate models returned the turtles and crocodiles grouping, whereas the GHOST model returned the turtles and archosaurs grouping. Therefore, the GHOST model is not misled by the saturation found at codon position 3 sites, whereas the discrete Γ and PDF rate models are.

**Figure 3:**
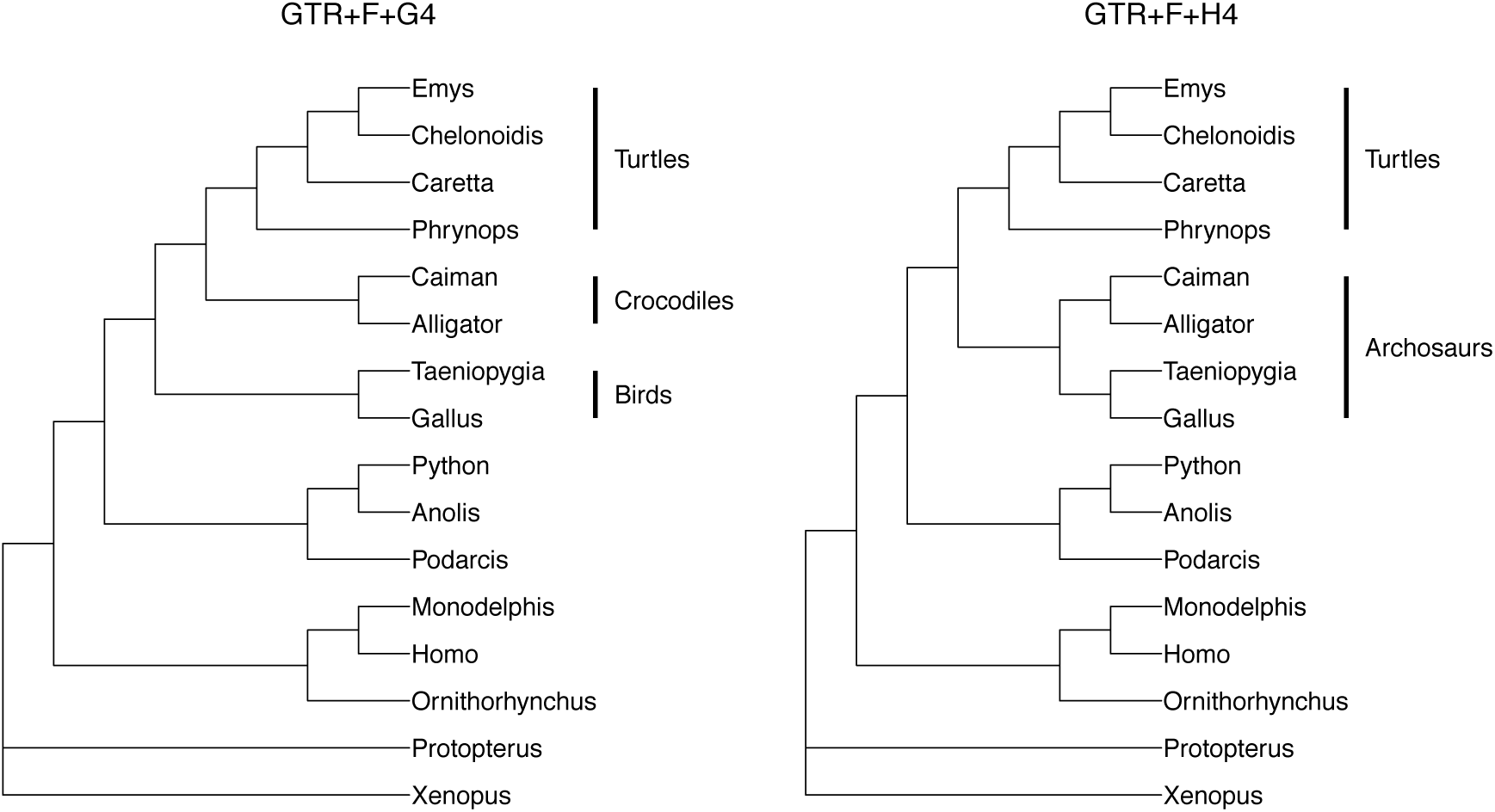
Turtle alignment - The two different topologies obtained from the turtle alignment. The topology on the left is returned by the 4-class discrete Γ and PDF rate models and places turtles as sister to crocodiles. The topology on the right is returned by the 4-class unlinked GHOST model and places turtles as sister to archosaurs (crocodiles and birds).

### Convergent Evolution of the Na_v_1.4a Gene Among Teleosts

#### Model selection and interpretation

To investigate its performance on empirical data, we applied the GHOST model to the coding region of a sodium channel gene, Na_*v*_1.4a, for 11 teleost species. Zakon et al. (2006) demonstrated the role of this gene in the convergent evolution of the electric organ amongst electric fish species from South America and Africa. AIC determined that GTR+FO*H4 (AIC=27602) provided the best fit between tree, model and data (Supplementary Fig. S4). Conversely, BIC determined that GTR+FO*H2 provided the best fit. Examining the class weights and trees (Figure 4) inferred by GTR+FO*H4 indicates that all classes have non-negligible weight (minimum class weight is 0.13) and all four trees appear reasonably distinct. Thus, we conclude that there are no obvious signs of overfitting present, and we accept four classes as optimal for this alignment. We also tested the empirical base frequencies version (GTR+F*H4, AIC=27749) and linked substitution rates version (GTR+FO+H4, AIC=27860). Each of these models returned a significantly higher AIC value, indicating that the unlinked version provided the best fit. We then tested the PDF rate model, finding that the best such model had six classes (GTR+FO+R6), but still a much higher AIC (27813) than that of the GTR+FO*H4 model. In order to confirm the stability of the parameter estimates we repeated the analysis using the GTR+FO*H4 model 100 times.

**Figure 4:**
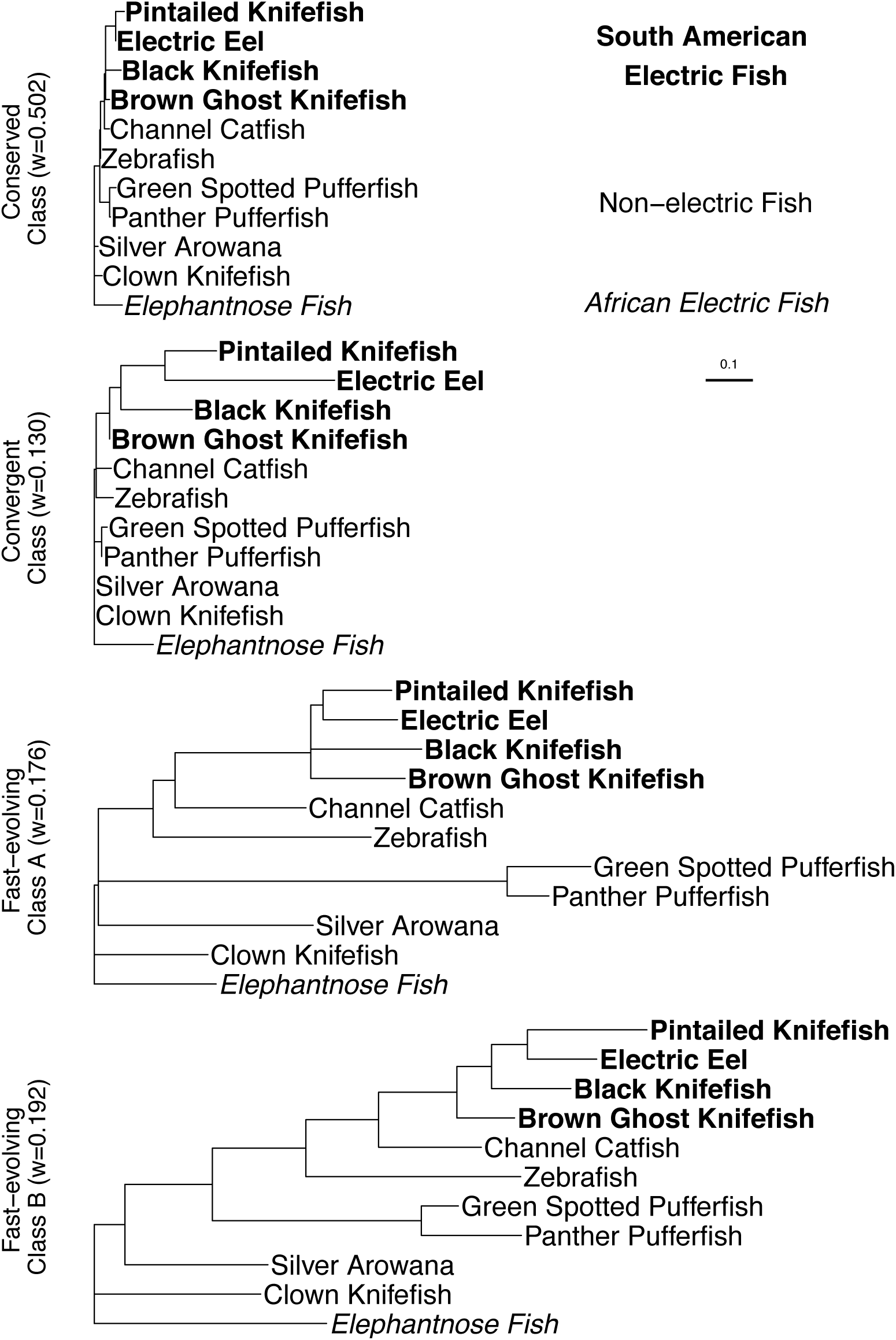
The four trees inferred under the General Time Reversible, four-class mixture model (GTR+FO*H4) for the electric fish data. The classes are displayed in order of increasing tree size, as determined by the sum of the branch lengths. We refer to this as the total tree length (TTL): TTL_*Cons*_ = 0.23, TTL_*Conv*_ = 0.99, TTL_*F EA*_ = 4.06 and TTL_*F EB*_ = 4.18.

We then partitioned the electric fish sequence alignment into three partitions, based on codon position (CP). PartitionFinder suggested GTR+FO+G4 (GTR with inferred equilibrium base frequencies plus discrete Γ with four classes) for both the CP1 and CP2 partitions, and GTR+FO+I+G4 (same as above but with the inclusion of an invariable sites class) for the CP3 partition. We used IQ-TREE to run the codon partition model with the models indicated by PartitionFinder. The trees inferred by the partition model can be found in Supplementary Figure S19.

#### Interpretation of results

We labelled the four classes inferred by IQ-TREE under the GTR+FO*H4 model in order of increasing TTL: the ‘Conserved Class’ (TTL_*Cons*_=0.23), the ‘Convergent Class’ (TTL_*Conv*_=0.99), ‘Fast-evolving Class A’ (TTL_*F EA*_=4.06) and ‘Fast-evolving Class B’ (TTL_*F EB*_=4.18). Of particular interest is the Convergent Class, so named as it corresponds well to Zakon *et al*. ‘s (2006) hypothesis of convergent evolution of Na_*v*_1.4a among the South American and African electric fish clades. They explained that the Na_*v*_1.4a gene arose from a single gene duplication event which occured in a species ancestral to all 11 fish species in the alignment, and was historically expressed in muscle tissue. They then show that the gene is now expressed in the electric organ of all but one of the electric fish species in both the South American and African electric fishes, but obviously not in the non-electric fishes. Since these lineages constitute two separate clades, one conclusion that can be drawn is that this morphological trait evolved twice independently, once in the South American clade and once in the African clade. Hence, this appears to be an interesting example of convergent evolution (convergent at the morphological level, but not necessarily at the molecular level). The inferred tree associated with the Convergent Class displays much more evolution in the electric rather than the non-electric fish lineages (Fig. 5). This is indicative of either a relaxation of purifying selection pressure, an introduction of positive selection pressure or a combination of both. The notable exception is the Brown Ghost Knifefish, which appears relatively conserved. The Brown Ghost Knifefish is unique amongst the electric fish in the dataset, in that its electric organ has evolved from neural rather than muscle tissue. Consequently, in the Brown Ghost Knifefish the *Na*_*v*_*1.4a* gene is still expressed in muscle, just as it is in the non-electric fish. The distinction in terminal branch length between the Brown Ghost Knifefish and the other electric fishes offers compelling evidence that the GHOST model has identified a subtle component of the historical signal related to the convergent evolution of *Na*_*v*_*1.4a*, as opposed to returning a somewhat arbitrary combination of numerical parameters that happen to maximize the likelihood function. To further verify that this conclusion was justified, we examined the trees inferred under the GTR+FO*H5 and GTR+FO*H6 models. If a convergent evolution signal is indeed present in the alignment then it should also be revealed under these models. Supplementary Figures S20 and S21 show the trees inferred by the five and six class model respectively. The third class in each Figure appears to capture a similar signal to that captured by the Convergent Class of the four class model. The ability of the GHOST model to isolate such a small component of the signal (the inferred weight of the convergent class being 0.13, the smallest of the 4 classes) is encouraging. Furthermore, we can hypothesize that the sites belonging with high probability to the Convergent Class are likely to have been influential in the functional development of the electric organ.

**Figure 5:**
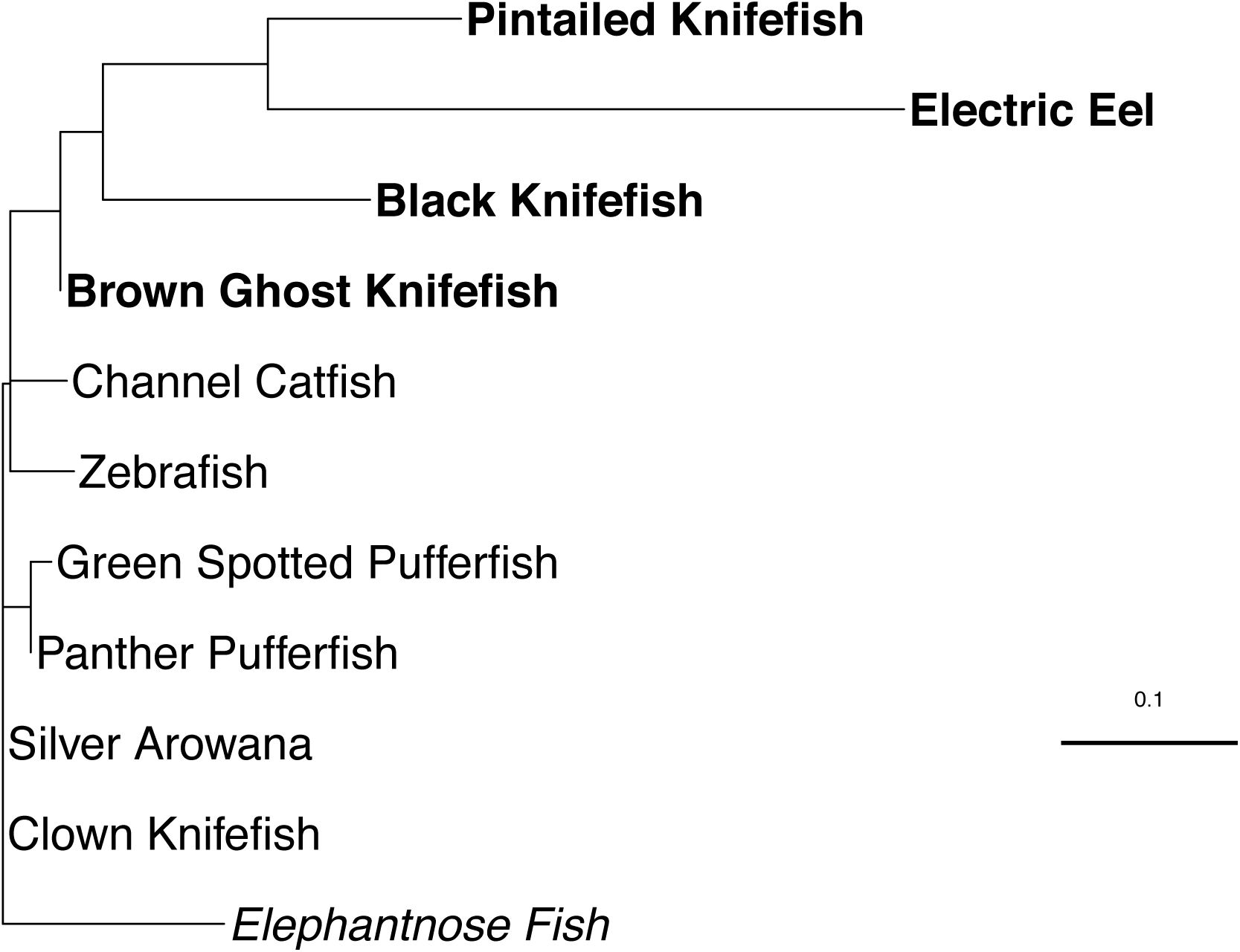
The convergent class inferred by ML-GTR+FO*H4. The 11 fish species comprised four South American electric fish (bold), one African electric fish (italics), and six non-electric fish (normal font) from various locations. The tree for this class shows that in comparison to the electric fish, the non-electric species are relatively conserved.

#### Soft classification of sites to classes

The soft classification of sites to classes facilitates the prospective identification of functionally important sites in an alignment. Zakon et al. (2006) report several amino-acid sites from the dataset that are influential in the inactivation of the sodium channel, a process critical to electric organ pulse duration. Figure 6a shows that these sites generally have a higher than average probability of belonging to the convergent class in at least one codon position. For example, at position 647, an otherwise conserved proline (codon CCN) is replaced by a valine (GTN) in the Pintailed Knifefish and a cysteine (TGY) in the Electric Eel. Unique substitutions at codon positions 1 and 2 are necessary for both of these amino-acid replacements, and we find these two sites have a very high probability of belonging to the convergent class. With this result in mind, for each amino acid we summed the probability of codon positions 1 and 2 belonging to the Convergent Class. Figure 6b shows the results for the eight amino-acid sites with the highest score. Comparing the magnitude of these bars with those of the amino-acid sites in Figure 6a (which are known to be functionally important), one is led to suspect that these amino acids might also be critical to the operation of the sodium channel gene. Given that there are many other sites in the alignment with a high probability of belonging to the convergent class, one can envisage the GHOST model helping to identify sites of potential functional importance in an alignment, thereby focusing the experimental work of biologists.

**Figure 6:**
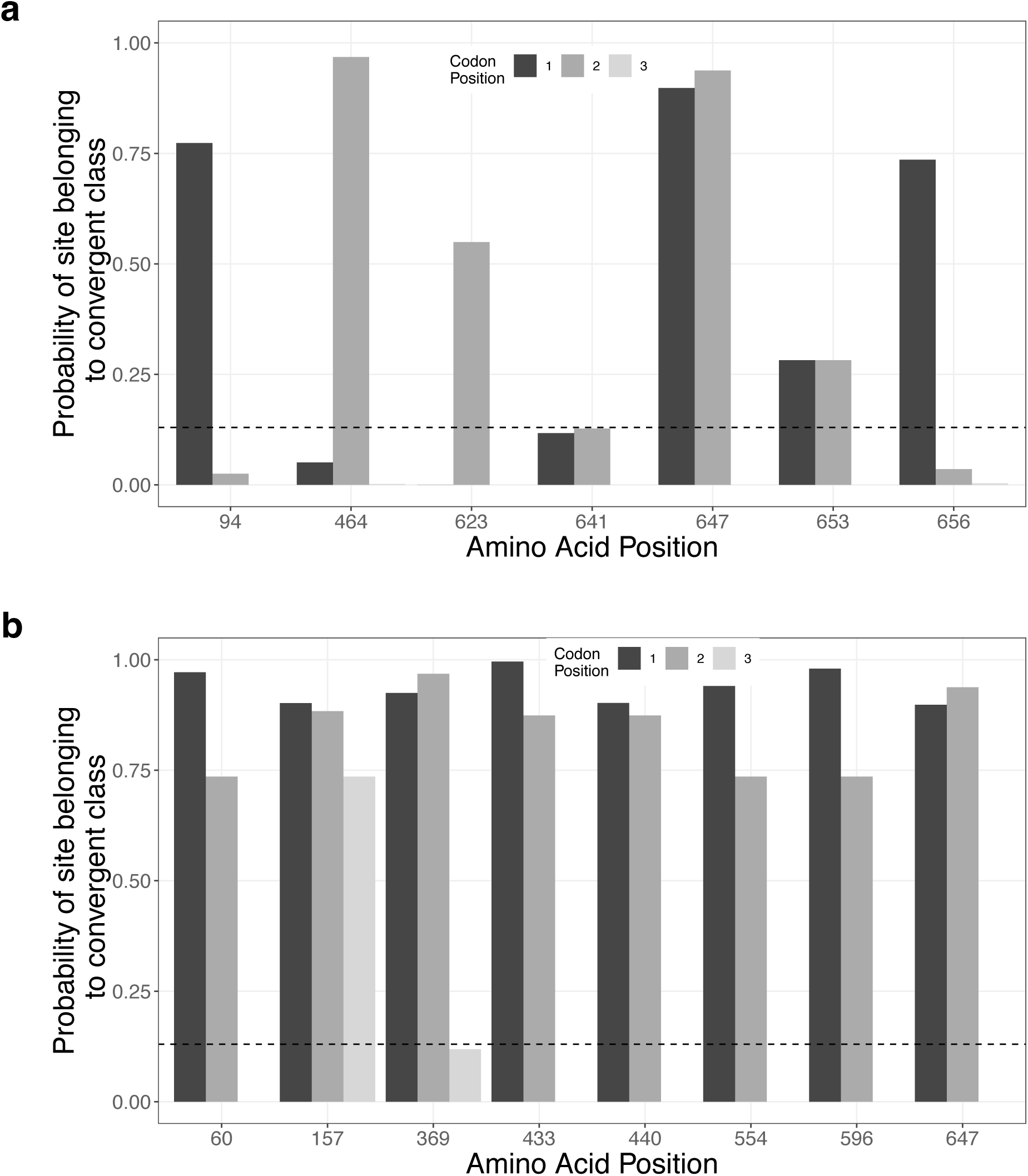
Probability of sites belonging to the convergent class by codon position. (a) The amino-acid positions selected correspond with those identified by Zakon et al. (2006) as being functionally important to the inactivation of the Na^+^ channel gene. The horizontal dotted line at 0.13 represents the average probability of belonging to the convergent class over all sites in the alignment. (b) The amino-acid positions selected correspond to those with the highest probability of belonging to the convergent class, summed across the first two codon positions.

In addition to providing insight on an individual site basis, the soft classification can also help to inform us about the nature of the classes themselves. Generally speaking, the branch lengths inferred by IQ-TREE can be interpreted as the expected number of substitutions per site. Therefore, summing the weighted TTLs for each of the inferred classes tells us that we expect 1.766 substitutions per site under the inferred model. Table 2 reports the contributions to this figure, stratified by codon position and class. If class membership and codon position were independent attributes of each site, then we should expect the contribution of each codon position to be approximately one third for each class. This is not what we observe. Overall we can see that sites in CP1(23%) and CP2 (16%) contribute only 39% of the total of 1.766 substitutions per site. However, within the Conserved and Convergent Classes, sites in CP1 and CP2 are responsible for 90% and 76% of their contribution respectively. This would suggest that a comparatively larger proportion of the substitutions attributed to these classes are non-synonymous: resulting in amino-acid replacements that influence the fitness of the organism. We can therefore conclude that even though the Conserved and Convergent Classes are smallest (as determined by substitutions per site), they appear to be the primary catalyst of evolution via natural selection within *Na*_*v*_*1.4a* amongst these species.

**Table 2:** Expected number of substitutions per site (bold) for the electric fish alignment, weighted by class and separated by codon position (CP). For each inferred class, the expected substitutions per site are calculated by multiplying the total tree length (TTL) by the class weight. The CP1, CP2 and CP3 columns show the contribution to these figures from only the sites within each CP. The grand total indicates that under the parameters inferred by ML-GTR+H4 we would expect 1.766 nucleotide substitutions per site. We can then see, for example, that the Convergent Class is responsible for 0.129 of these substitutions per site. Finally, of the 0.129 substitutions per site attributable to the Convergent Class, 0.051 (or 40%) is the contribution from sites in CP1, 0.047 (36%) is the contribution from sites in CP2 and 0.031 (24%) is the contribution from sites in CP3.

| Class | CP1 | CP2 | CP3 | Subs/site |
| --- | --- | --- | --- | --- |
| Conserved | 0.049 (41%) | 0.058 (49%) | 0.012 (10%) | **0.119** |
| Convergent | 0.051 (40%) | 0.047 (36%) | 0.031 (24%) | **0.129** |
| Fast-evolving A | 0.135 (19%) | 0.076 (11%) | 0.504 (70%) | **0.715** |
| Fast-evolving B | 0.175 (22%) | 0.100 (12%) | 0.528 (66%) | **0.803** |
| All Classes | 0.410 (23%) | 0.280 (16%) | 1.076 (61%) | **1.766** |

#### Comparison to the Partition Model

It is apparent upon examination of the trees in Supplementary Figure S19 that the evidence of convergent evolution highlighted by the GHOST model (Fig. 5) has not been recovered by the codon-based partition model. None of the three trees in Supplementary Figure S19 have the distinctive pattern, whereby the majority of the total tree length is associated with the electric fish species (with the exception of the Brown Ghost Knifefish). The reason that the partition model was unable to recover this signal has to do with the relative contribution of sites from each CP to the Convergent Class. Table 2 indicates the extent to which the substitutions associated with the Convergent Class are attributable to CP1 sites (40%), CP2 sites (36%) and CP3 sites (24%). The partition model constrains the analysis, such that sites in different CPs are modeled independent of each other. It is impossible for a model constrained in such a way to effectively recover the convergent evolution signal because the signal is distributed across all three partitions. The decision to partition the data based on codon position may make sense superficially, but in doing so the analysis is constrained and the results are compromised. We no longer have the ability to uncover the evolutionary stories concealed within the data. We can only hope to obtain those stories that happen not to conflict with the assumptions and constraints that have been placed on the analysis *a priori*. Minimizing these assumptions and constraints where possible, while computationally expensive, is necessary in order to illuminate the evolutionary history without distorting it in the process.

### On the Identifiability of the GHOST Model

An ongoing concern regarding parameter-rich mixture models has been whether or not they are identifiable. There are several examples of theoretically non-identifiable mixture models in the literature (Matsen and Steel, 2007;Štefankovič and Vigoda, 2007b). These examples have inspired much theoretical work on the identifiability or otherwise of different types of phylogenetic mixture models (Allman and Rhodes, 2006;Štefankovič and Vigoda, 2007a; Allman et al., 2008; Allman and Rhodes, 2008; Steel, 2010; Allman et al., 2011). Of particular interest to the current study, Allman et al. (2011) showed that for a single topology, four taxa, two-class mixture under the JC model, only the tree topology is identifiable but not the branch lengths. This provides a theoretical justification for the procedure carried out by K&T (and replicated here), measuring performance of the models based only on recovery of the topology and paying no attention to recovery of branch length parameters. With regard to the identifiability of the GHOST model more generally, we rely on a result from Rhodes and Sullivant (2012). They established an upper bound on the number of classes, *m*, for which tree topology, branch lengths and model parameters are identifiable, as a function of the number of character states, *κ*, and the number of taxa, *n*:

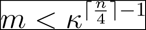

For the simulations we carry out in the current study, with 12 taxa and four character states, the model is identifiable up to a maximum of 16 classes. For 32 taxa and four character states, the model is identifiable up to a maximum of 16,384 classes. In the case of the electric fish dataset, with four character states and only 11 taxa, the model is identifiable up to 16 classes. However, there is a technical caveat. The result is shown based on assuming a general Markov model across the tree. There are specific choices of parameters that can result in non-identifiability, but these are of little concern in practical data analysis. Problems arise only when the parameters selected collapse the parameter space to some lower dimension. For example, we could fit the GTR model but if we chose parameters such that all base frequencies were equal and all substitution rates were equal then we are in fact using a JC model, and identifiability may be compromised. However, these technical examples of non-identifiability are not relevant in practice, as in the absence of any constraints there is no reasonable chance of inferring parameters that collapse the parameter space in such a way.

## Conclusion

Heterotachy has been somewhat of an Achilles heel for ML since K&T published their study. The ML implementation of the GHOST model in IQ-TREE represents a positive advance for ML-based phylogenetic inference. Through minimization of model assumptions, the GHOST model offers significant advantages and flexibility to infer heterotachous evolutionary processes, illuminating historical signals that might otherwise remain hidden. Owing to the diversity of selective pressures acting on different genes, the GHOST model seems well suited to the analysis of phylogenomic datasets (albeit with the limitation of being constrained to a single tree topology), commonly used to address deep phylogenetic questions. Forthcoming empirical studies will further compare the performance of the GHOST model to currently popular phylogenomic analysis tools, such as partition and CAT models. In addition, further simulation studies will help to better establish practical limitations to the use of the GHOST model, in terms of number of taxa, number of classes, sequence length and computation time. Many opportunities for refining or extending the GHOST model also present themselves. It could be used in conjunction with partition models, to account for heterotachy within partitions; if the data suggests correlation of some branch lengths across classes, then these could be linked to decrease the parameter space; it could form the basis of a test for heterotachy itself, by comparing results obtained under the GHOST model to those obtained under a discrete Γ or PDF rate model. One can also envisage many other potential applications for the GHOST model. It may be insightful when applied to datasets for which the topology is poorly supported or disputed. It could also provide more accurate parameter estimates, leading to sounder divergence date estimation. The model provides intuitive, biologically meaningful visualizations of the different evolutionary pressures that act on a group of taxa. Structural biologists may find it useful for highlighting functionally important areas within proteins. We have demonstrated its use as a method for identifying changes in selection pressure, as well as bringing to light evidence of convergent evolution. Similarly, one can envisage the GHOST model illuminating the subtle evolutionary relationships between hosts and parasites, disease and immune cells, or the countless evolutionary arms races that are observed throughout the natural world.

## SUPPLEMENTARY MATERIAL

Supplementary material, including further simulation results, figures and IQ-TREE command line instructions can be found in the Dryad data repository (doi TBA).

## ACKNOWLEDGEMENTS

The authors would like to thank Elizabeth Allman, John Rhodes and Edward Susko for helpful discussions about the manuscript.

B.Q.M. and A.v.H were supported by the Austrian Science Fund (FWF I-2805-B29).

## COMPETING FINANCIAL INTERESTS

The authors declare no competing financial interests.

