## Supplementary Material for "GHOST: Recovering Historical Signal from Heterotachously-evolved Sequence Alignments"

#### *Kolaczkowski & Thornton simulations*

We followed the simulations of Kolaczkowski and Thornton (2004) precisely and compared the performance of MP, ML-JC (ML under a JC model) and ML-JC+H2 (ML under JC with 2 GHOST classes). We used *Seq-Gen* (Rambaut and Grassly, 1997) to simulate nucleotide sequences on two symmetric, 4-taxon trees of identical topology (see Supplementary Fig. S1a) using the JC (Jukes and Cantor, 1969) model of evolution. The branch lengths were constructed such that each tree comprised of two non-sister long branches (length $p$) and two non-sister short branches (length $q$) separated by an internal branch (length $r$). We replicated three separate experiments previously carried out by K&T.

*Experiment 1.*— We fixed $p = 0.75$ and $q = 0.05$ (see Supplementary Fig. S1a) and varied the internal branch length, $r$, on the interval $[0.01, 0.4]$ in increments of 0.01. For each value of $r$, 200 simulated MSAs were constructed by concatenating two sub-alignments of equal length, one simulated on each of the trees in Supplementary Figure S1a. We carried out phylogenetic inference on each MSA using MP, ML-JC and ML-JC+H2. The experiment was repeated for sequence lengths of 1,000, 10,000 and 100,000 bp. The results are shown in Supplementary

Figure S1b. We found that both ML-JC and MP were misled when $r$ was short, but as $r$ increased the performance of MP recovered before ML. For a sequence length of 100,000 bp, MP was misled to some extent for $r < 0.24$ and ML-JC was misled for $r < 0.3$. These findings mirrored those of K&T precisely. However, the ML-JC+H2 model was never misled. Supplementary Figure S1b shows that given sufficient sequence length, the ML-JC+H2 model inferred the correct topology from the heterogeneous sequences 100% of the time with $r$ as low as 0.01. These results demonstrate that the ML-JC+H2 model can correctly infer the tree topology when both ML-JC and MP are misled by the heterotachous nature of the data.

*Experiment 2.*— We tested nine different combinations of $p \in \{0.3, 0.5, 0.7\}$ and $q \in \{0.001, 0.1, 0.2, 0.3, 0.4\}$ (see Supplementary Fig. S1a). For each of the three methods/models (MP, ML-JC and ML-JC+H2) and at each combination of $p$ and $q$ we determined the smallest value of $r$ (subject to the minimum $r = 0.001$), denoted $BL_{50}$ by K&T, such that the correct topology was returned at least 50% of the time for simulated alignments of length 10,000 bp. The results (Supplementary Fig. S2) indicate that ML-JC+H2 consistently outperformed the two alternatives, with the difference most apparent when the influence of heterotachy was strongest (most notably when $p$ is large and $q$ is small). Again, the results we observed for MP and ML-JC closely emulated the findings of K&T.

*Experiment 3.*— We tested the impact of varying the weight, $w$, of each of the two classes in the simulated MSAs for a variety of branch length combinations. Initially, $p$ and $q$ (see Supplementary Fig. S1a) were fixed at 0.75 and 0.05 respectively, with $r \in \{0.05, 0.15, 0.25\}$ and $w_1 \in$ {0.01, 0.1, 0.2, 0.3, 0.4, 0.5, 0.6, 0.7, 0.8, 0.9, 0.99}. The process was then repeated, this time with $p$ and $r$ fixed at 0.75 and 0.15 respectively, with $q \in \{0.05, 0.15, 0.25\}$ and $w$ as before. Sequence length was held fixed throughout at 100,000 bp (mirroring the experiment of K&T) and 200 replicates were simulated at each combination of branch lengths and weight. We found that for almost all branch length combinations ML-JC+H2 was able to recover the correct topology for all replicates. In the entire experiment, only one dataset (out of 13,200) returned the incorrect topology. The results of K&T indicate that ML-JC could not reliably recover the correct topology for all weights for any of the branch length combinations.

The positive performance of the GHOST model across the three K&T experiments should be expected in some sense, as ML-JC+H2 enjoys substantial advantage over the two alternatives. It is in no way misspecified, having the freedom to fit two classes evolved under the JC substitution model, precisely the conditions used to simulate the data. Conversely, ML-JC has only a single class and therefore is subject to model misspecification. No single set of branch lengths can reproduce the signal present in the simulated alignments. MP is obviously not subject to model misspecification as the method is non-parametric, but it is subject to the long-established artefact of long branch attraction (LBA) (Felsenstein, 1978). Felsenstein showed that having long non-sister branches separated by a relatively short internal branch can result in MP incorrectly inferring the long branches as sisters. Supplementary Figure S1a shows the two trees used for the classes in the mixture, both sharing the same AB|CD topology. The Class 1 tree has long terminal branches on the A and C lineages, therefore the LBA artefact biases MP towards the AC|BD topology. The Class 2 tree is in a sense the symmetric opposite of the Class 1 tree, it has long terminal branches on the B and D lineages so the result is the same: LBA biases MP towards the AC|BD topology.

Therefore, the successful replication of the K&T simulations is a necessary but not sufficient condition for the GHOST model's endorsement. It indicates that the ML implementation of the GHOST model within IQ-TREE's algorithm structure has been successful, but these simulations are on only four taxa and use the most simple model of sequence evolution. Moreover, they only focus on recovering the correct tree topology and not inferring branch length parameters.

#### *Software*

IQ-TREE can perform inference with the GHOST model on both nucleotide and amino-acid sequences, although it should be noted that simulation studies have only been carried out for nucleotide sequences. The GHOST model is executed in IQ-TREE v1.6 by augmenting the model argument as shown below. If one wants to fit a four-class, fully linked (model parameters are common to all classes) GHOST model with a GTR model of evolution, to sequences contained in `data.fst`, one would use the following command:

```
iqtree -s data.fst -m GTR+H4
```

The above command infers four sets of branch lengths, a single set of relative substitution rate parameters (which is common to all classes) and the weights of each class. The base frequencies are not inferred, but are taken from the empirical values observed in the alignment. So in effect the four classes only differ in that they each have their own set of branch lengths. However, we can gradually increase the complexity of the model if we so choose. To infer equilibrium base frequencies using maximum likelihood, instead of using the empirical base frequencies from the alignment, we add the `+FO` option:

```
iqtree -s data.fst -m GTR+FO+H4
```

The relative rate and base frequency parameters are still fully linked across all four classes. If one also wishes to infer separate GTR rate parameters and base frequencies for each class then the unlinked version is required:

```
iqtree -s data.fst -m GTR+FO*H4
```

This is the most general, fully unlinked version of the GHOST model. If one wishes to obtain a file with the probability of each site belonging to each class, then this can be done by using the `-wspm` option, as in:

```
iqtree -s data.fst -m GTR+FO*H4 -wspm
```

### Supplementary Figures and Tables

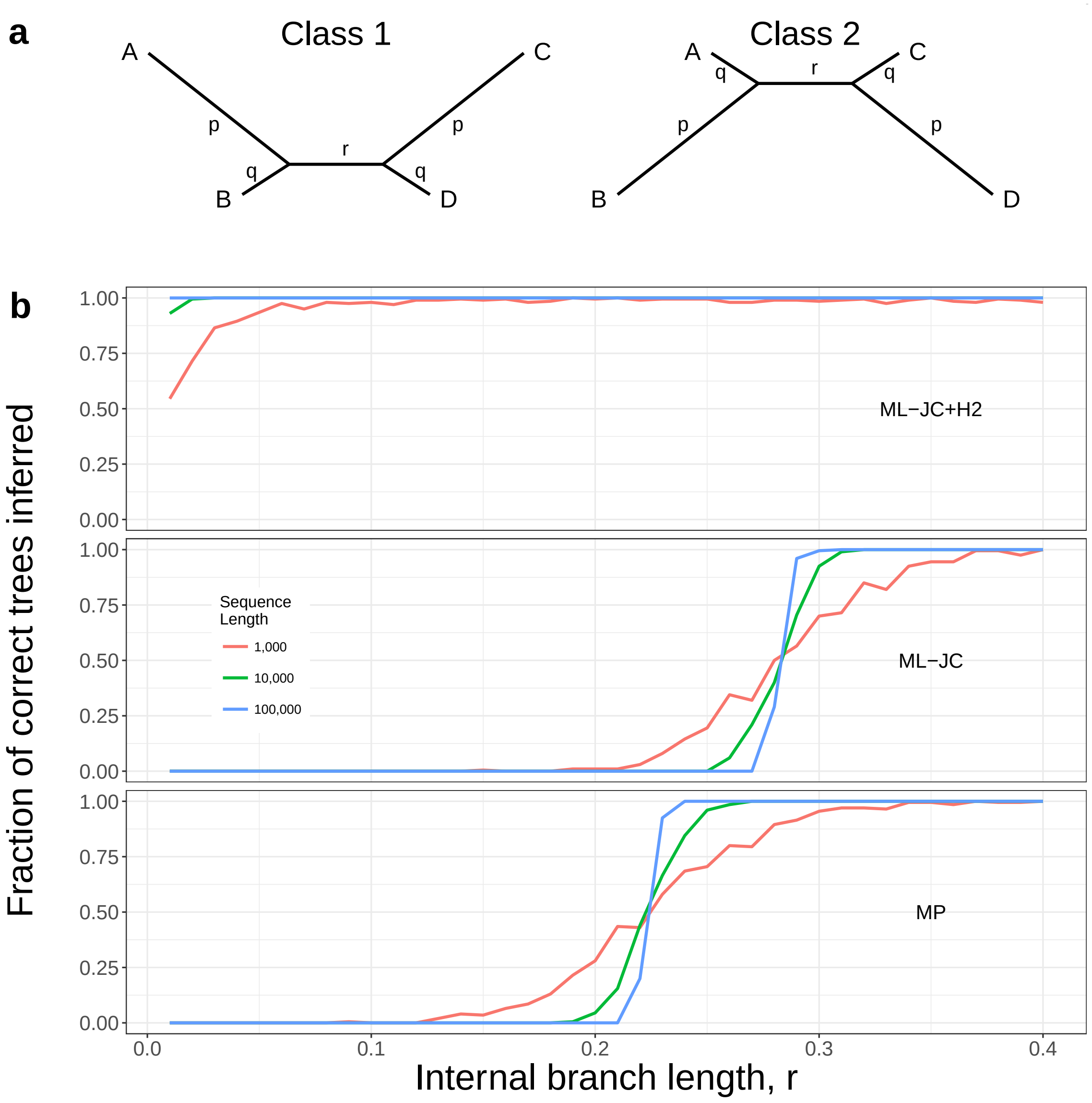


Figure S1: Replication of the simulations of Kolaczkowski & Thornton. (a) We simulated DNA sequences on two symmetric, 4-taxon trees of identical topology using the Jukes-Cantor (JC, 1969) model of nucleotide substitution. The branch lengths were constructed such that each tree comprised of two non-sister long branches and two non-sister short branches. Thus each tree was susceptible to long branch attraction (Felsenstein, 1978) (LBA). Importantly, the LBA artefact in both trees was complementary - the bias was in the direction of the AC|BD tree. (b) Performance of maximum likelihood (ML) using a JC, two-class mixture model (ML-JC+H2), ML using a single-class JC model (ML-JC) and maximum parsimony (MP) for data generated under strong heterotachy, $p$=0.75 and $q$=0.05. The length of the internal branch, $r$, was varied between 0.01 and 0.4 with 200 replicates at each value of $r$. ML-JC+H2 was able to reliably recover the tree topology for these data even when the internal branch is very short.

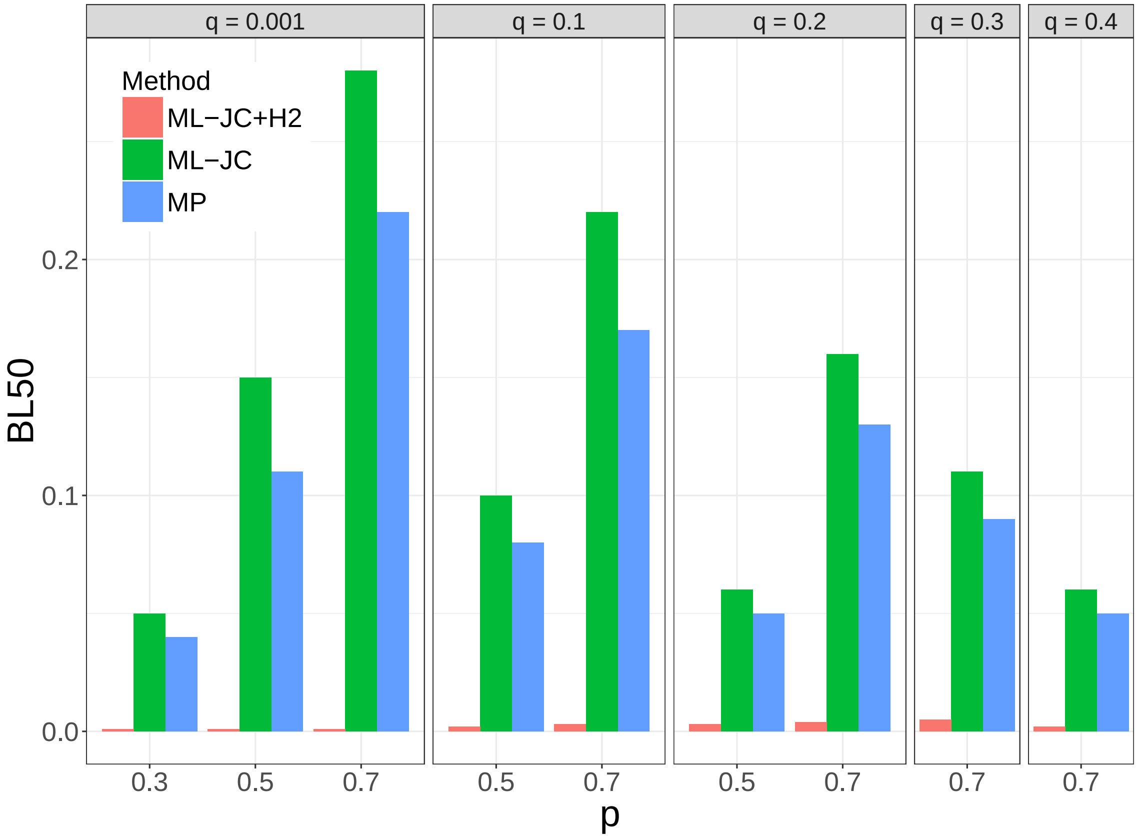


Figure S2: The ML-JC+H2 model outperforms MP and ML-JC over the range of heterotachous conditions tested by K&T, using 10,000 bp simulated alignments. They introduced the $BL_{50}$ measure as the minimum internal branch length required for the method to recover the correct tree topology at least 50% of the time. Small values of $BL_{50}$ indicate that the model is less likely to be misled by the heterotachous nature of the data.

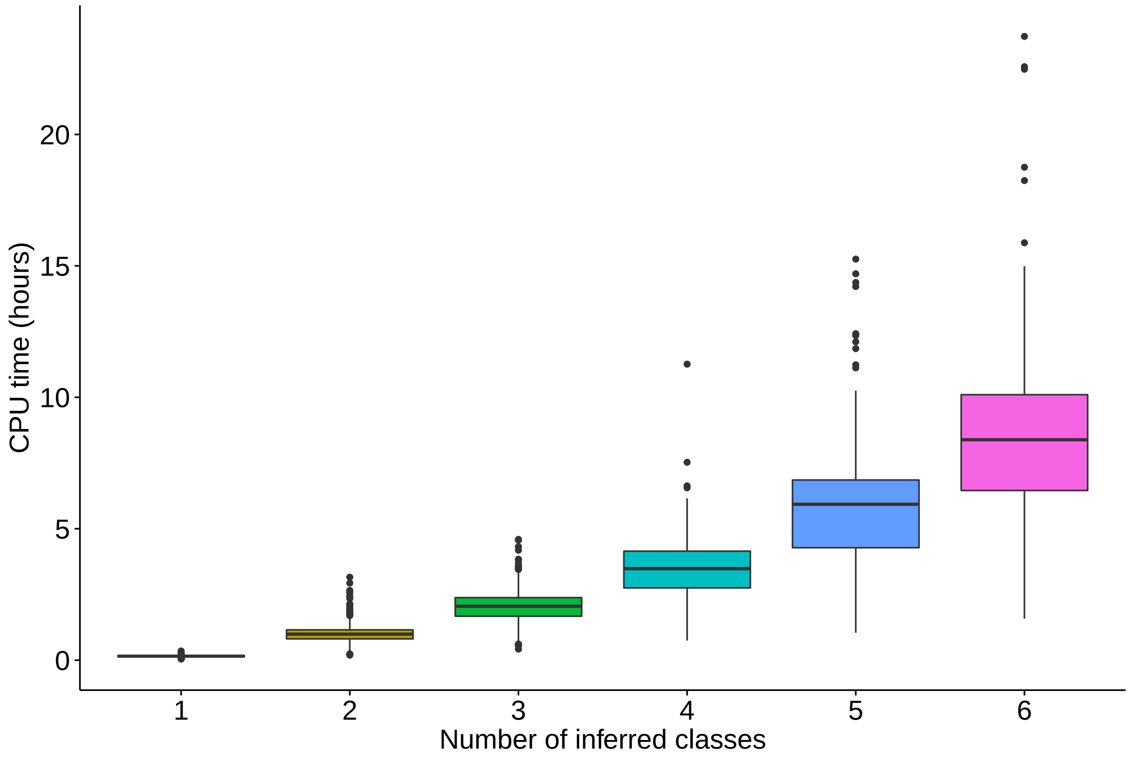


Figure S3: 32-taxon simulations, boxplots showing the computation time required for IQ-TREE to carry out ML inference under the GHOST model, as a function of the number of classes inferred.

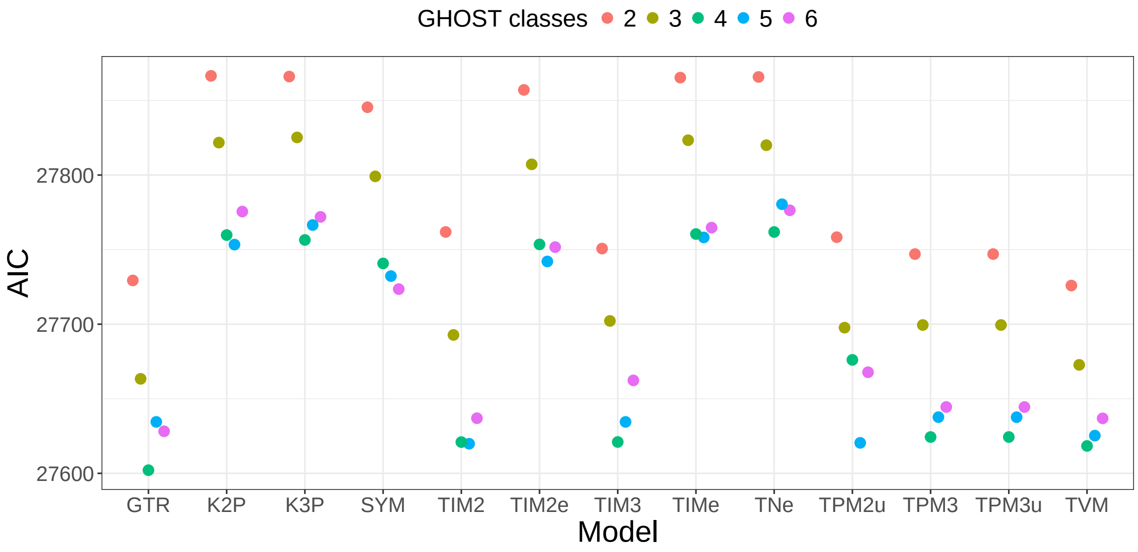


Figure S4: Results of model test procedure to select best substitution model and number of classes for the electric fish alignment, using AIC as the discriminating criterion. A total of 13 different nucleotide substitution models were tested, shown on the x-axis. Five separate IQ-TREE runs were carried out for each model, with the number of classes varied between two and six. GTR+FO*H4 was found to provide the best fit to the data.

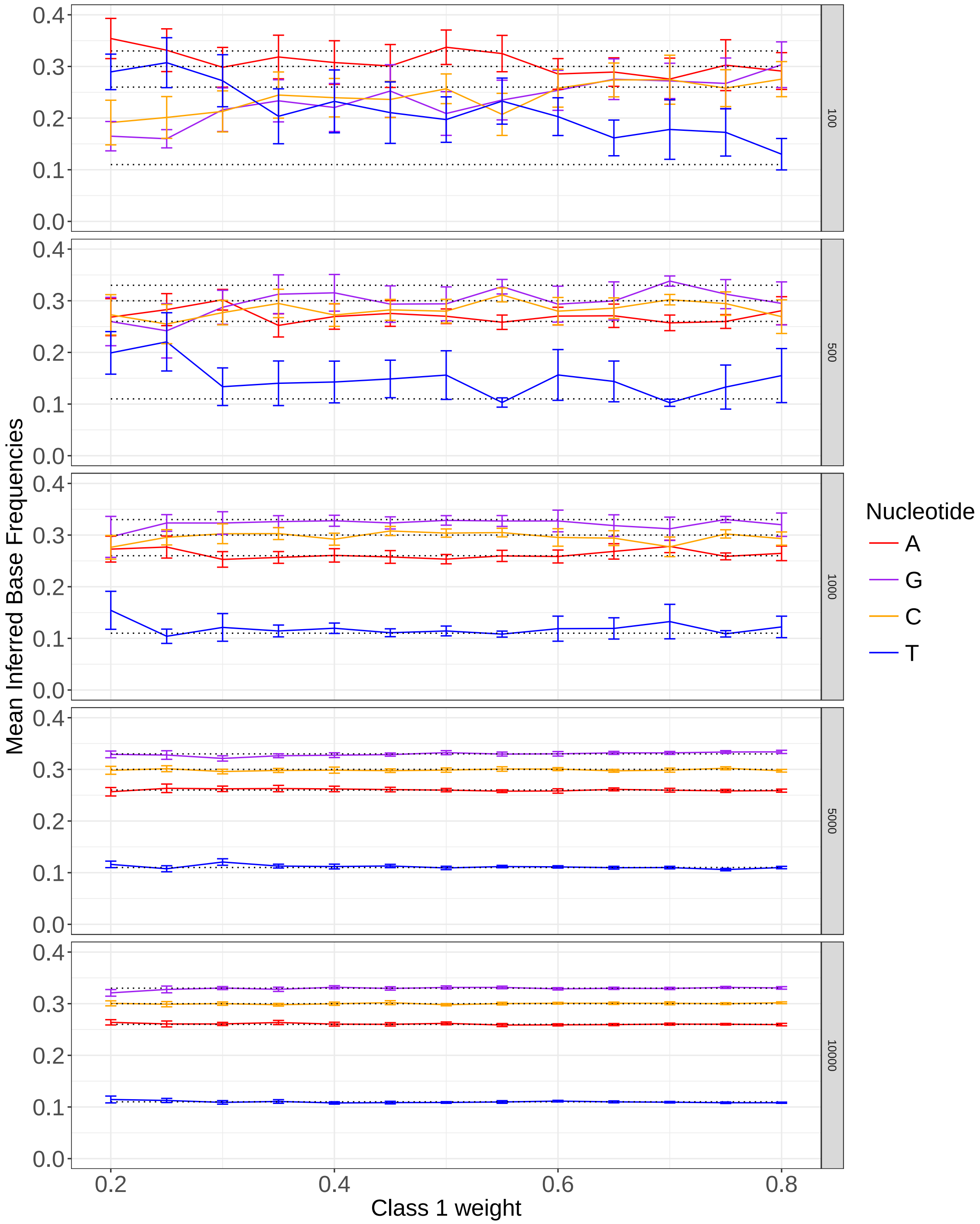


Figure S5: 12-taxon simulations - Class 1 inferred base frequencies vs Class 1 weight, faceted by sequence length. The sequence length is given in the right hand margin of each plot. The data points indicate the mean value of the base frequencies over the 20 simulated alignments at each value of Class 1 weight. The error bars represent $\pm 2$ standard errors of the mean. The dotted lines represent the true base frequency values used for data simulation.

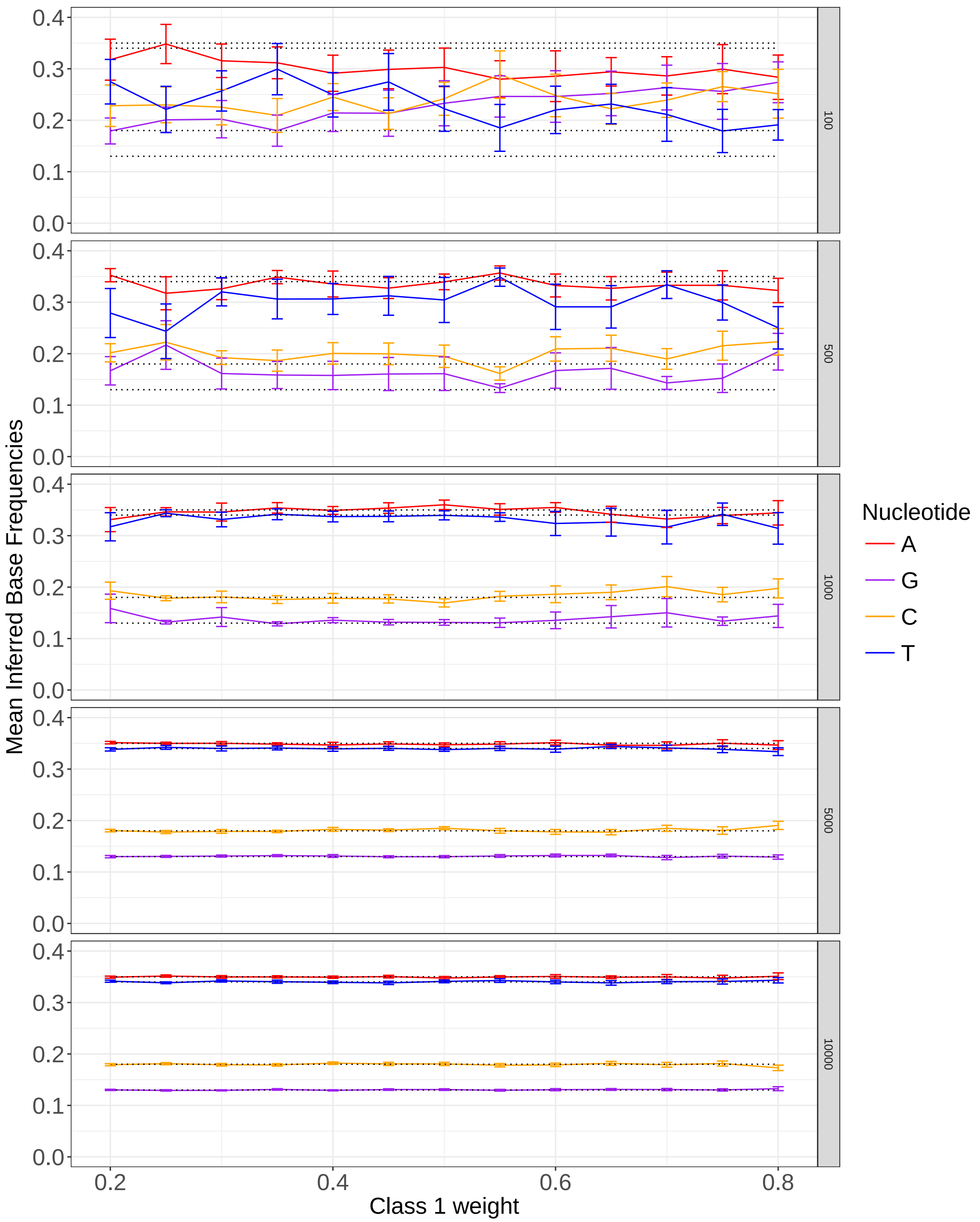


Figure S6: 12-taxon simulations - Class 2 inferred base frequencies vs Class 1 weight, faceted by sequence length. The sequence length is given in the right hand margin of each plot. The data points indicate the mean value of the base frequencies over the 20 simulated alignments at each value of Class 1 weight. The error bars represent ±2 standard errors of the mean. The dotted lines represent the true base frequency values used for data simulation.

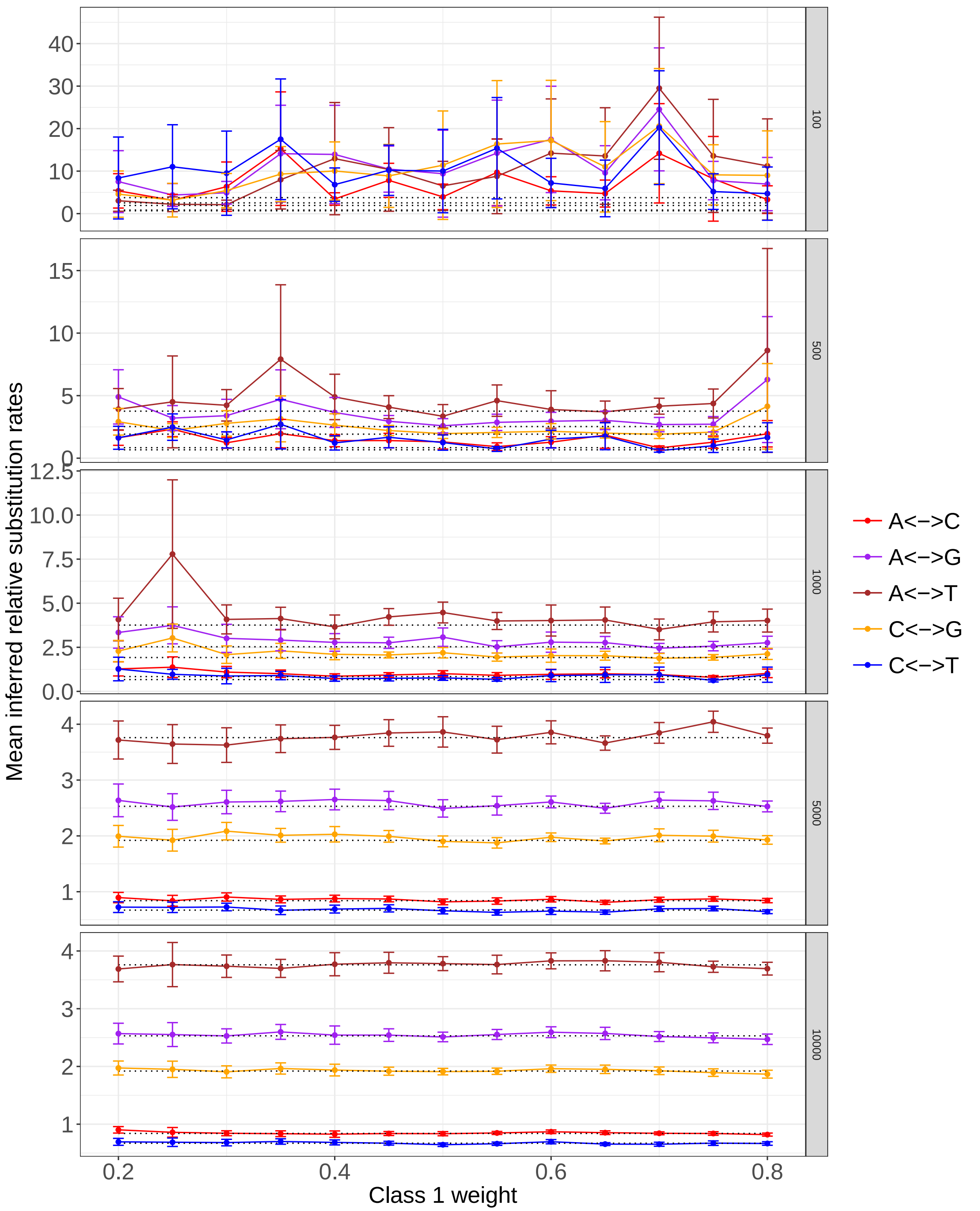


Figure S7: 12-taxon simulations - Class 1 inferred relative rates vs Class 1 weight, faceted by sequence length. The sequence length is given in the right hand margin of each plot. The data points indicate the mean value of the relative substitution rates over the 20 simulated alignments at each value of Class 1 weight, the error bars represent $\pm 2$ standard errors of the mean. The dotted lines represent the true relative substitution rate values used for data simulation.

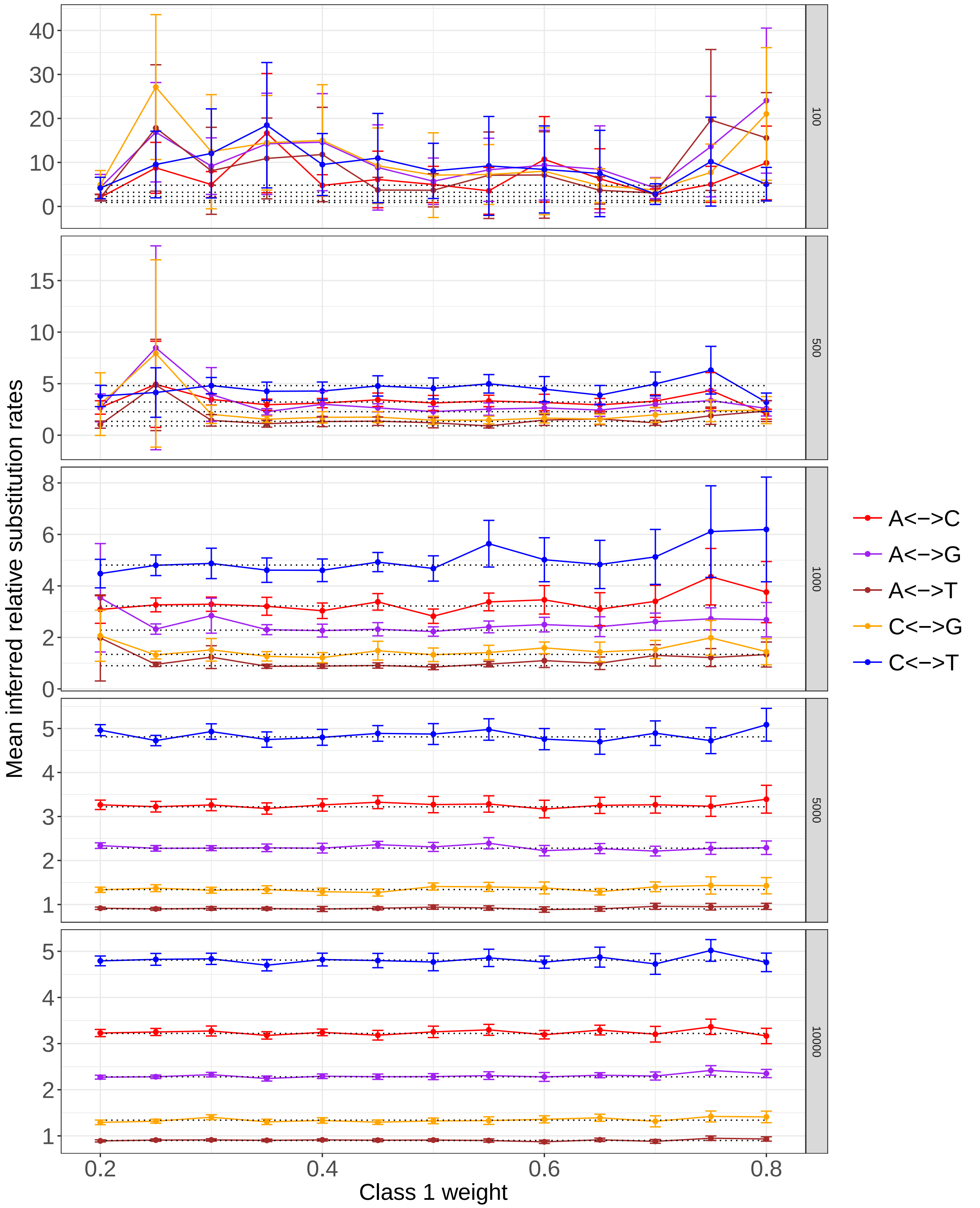


Figure S8: 12-taxon simulations - Class 2 inferred relative rates vs Class 1 weight, faceted by sequence length. The sequence length is given in the right hand margin of each plot. The data points indicate the mean value of the relative substitution rates over the 20 simulated alignments at each value of Class 1 weight, the error bars represent $\pm 2$ standard errors of the mean. The dotted lines represent the true relative substitution rate values used for data simulation.

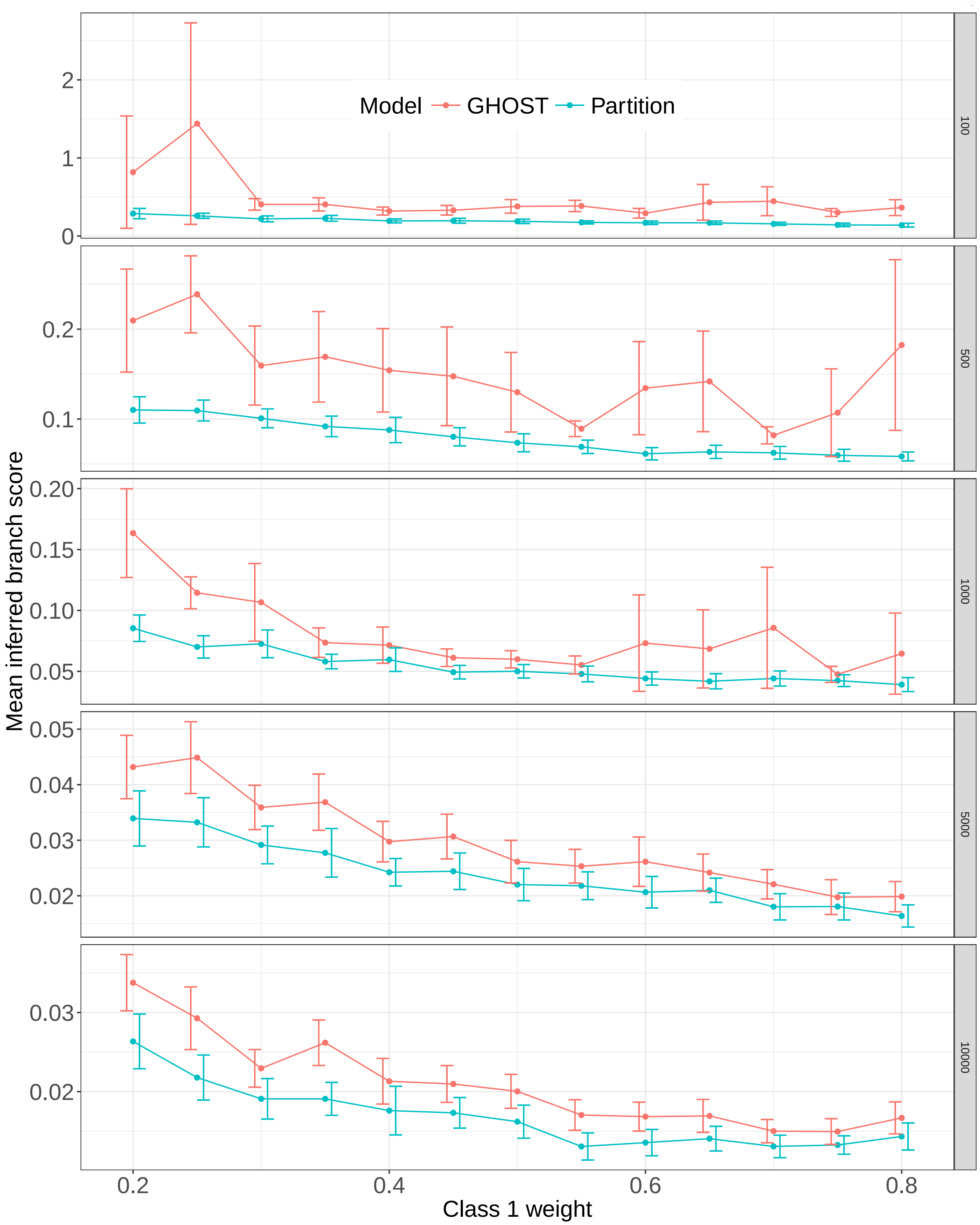


Figure S9: 12-taxon simulations - Class 1 branch score vs Class 1 weight, faceted by sequence length. The sequence length is given in the right hand margin of each plot. The data points indicate the mean value of the branch score over the 20 simulated alignments at each value of Class 1 weight. The error bars represent ±2 standard errors of the mean. Results obtained by under the GTR+FO*H2 are shown in red, those obtained under the partition model are shown in blue.

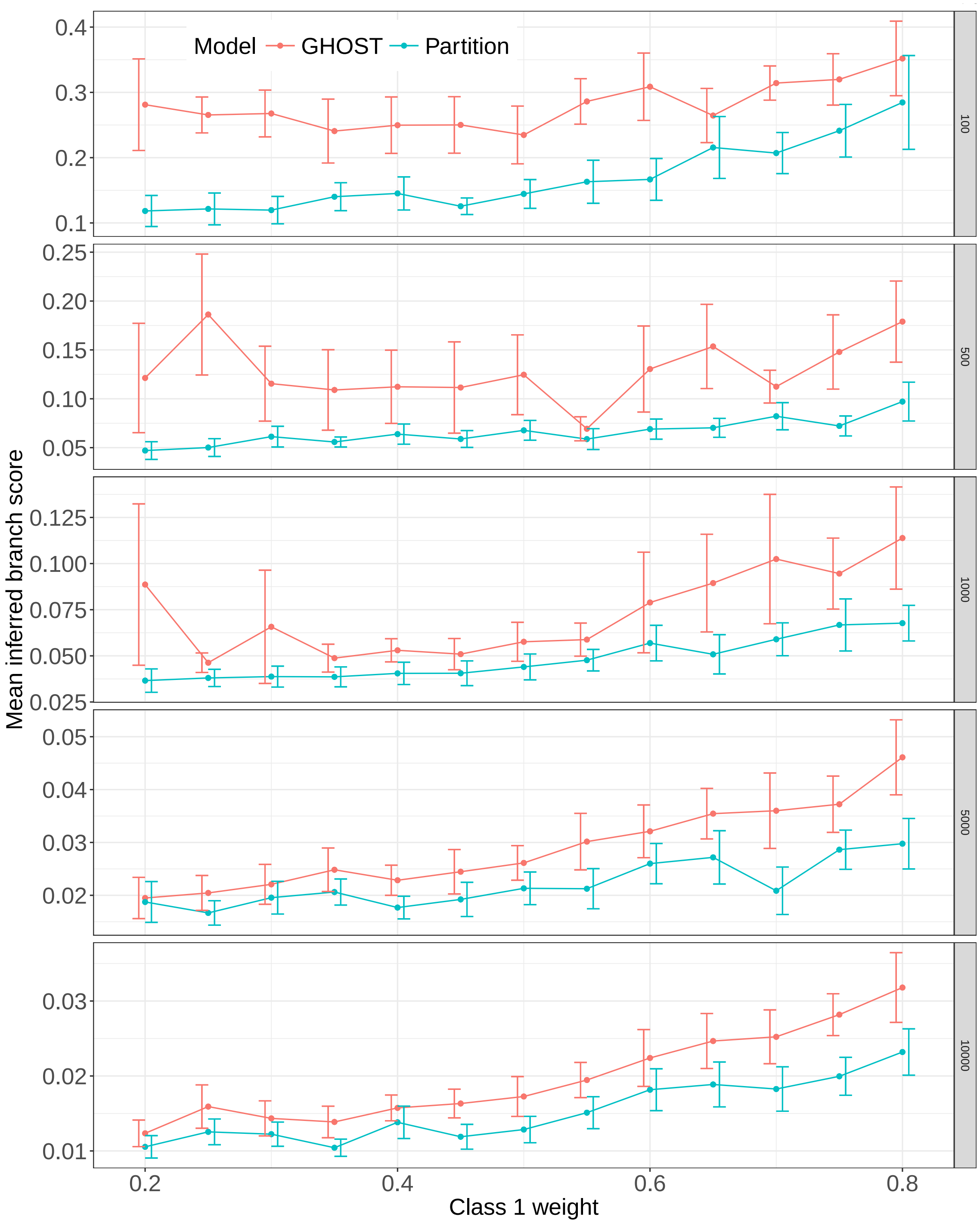


Figure S10: 12-taxon simulations - Class 2 branch score vs Class 1 weight, faceted by sequence length. The sequence length is given in the right hand margin of each plot. The data points indicate the mean value of the branch score over the 20 simulated alignments at each value of Class 1 weight. The error bars represent ±2 standard errors of the mean. Results obtained by under the GTR+FO*H2 are shown in red, those obtained under the partition model are shown in blue.

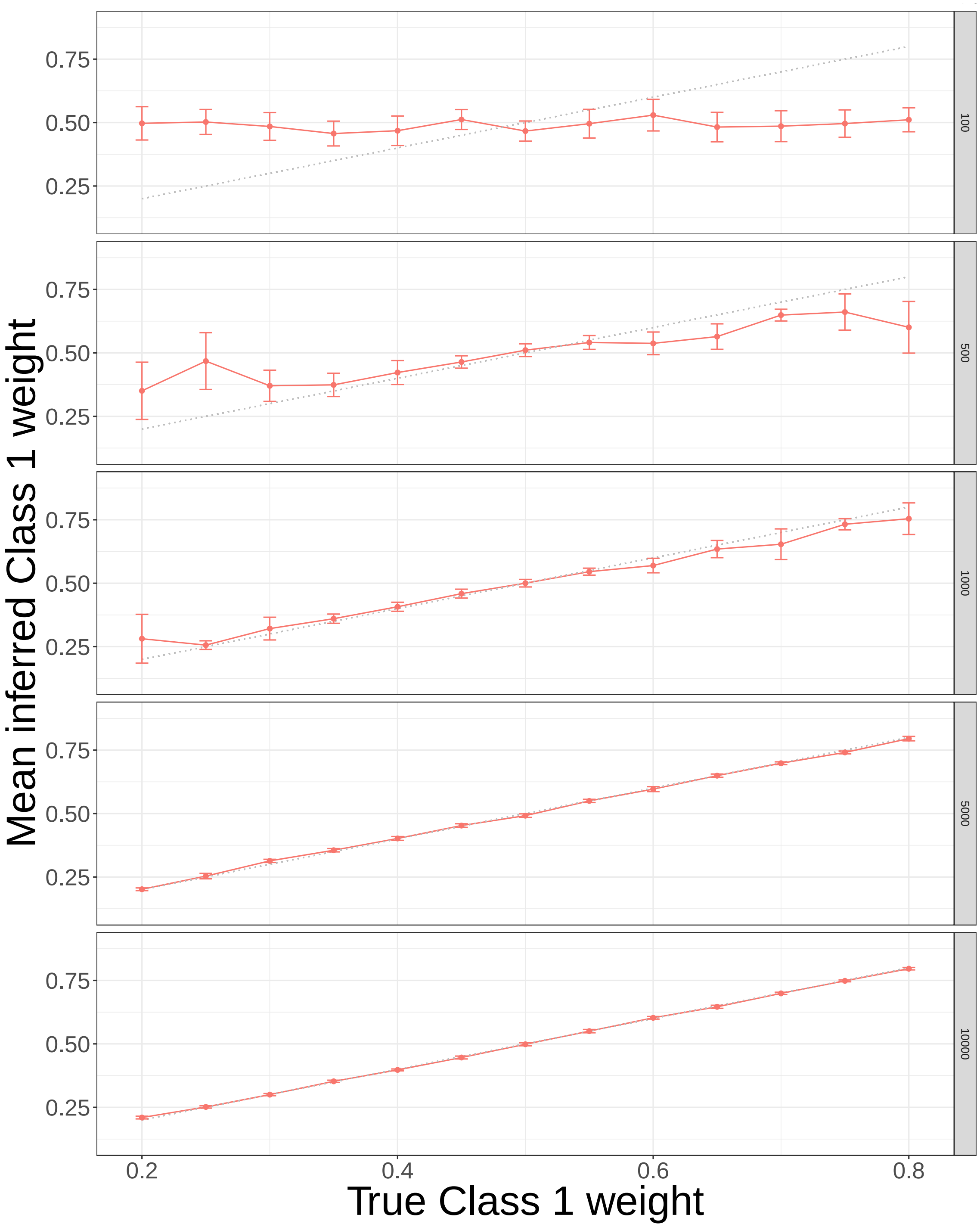


Figure S11: 12-taxon simulations - Class 1 inferred weight vs Class 1 weight, faceted by sequence length. The sequence length is given in the right hand margin of each plot. The data points indicate the mean value of the base frequencies over the 20 simulated alignments at each value of Class 1 weight. The error bars represent ±2 standard errors of the mean. The dotted lines represent the true Class 1 weight values used for data simulation.

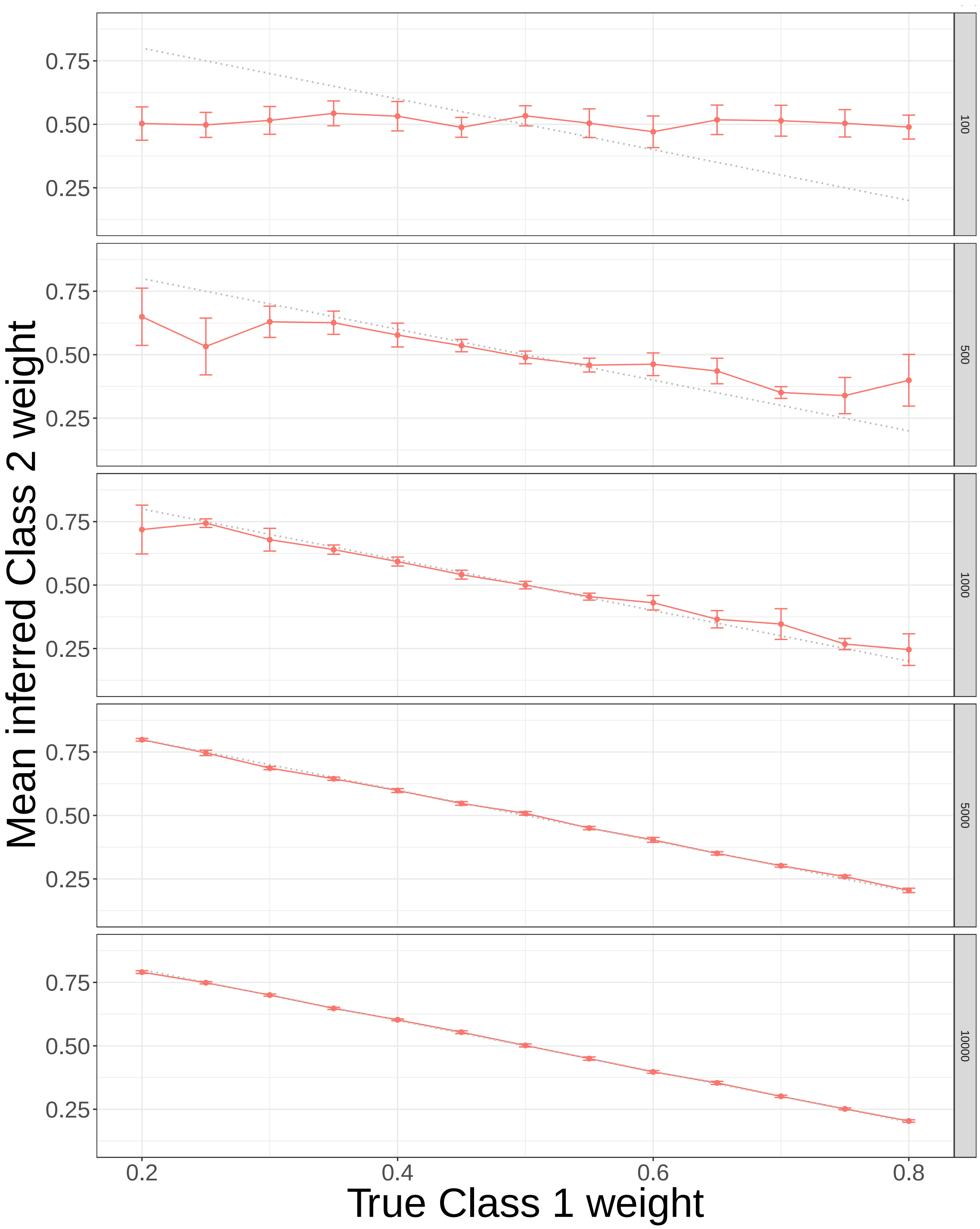


Figure S12: 12-taxon simulations - Class 2 inferred weight vs Class 1 weight, faceted by sequence length. The sequence length is given in the right hand margin of each plot. The data points indicate the mean value of the base frequencies over the 20 simulated alignments at each value of Class 1 weight. The error bars represent $\pm 2$ standard errors of the mean. The dotted lines represent the true Class 2 weight values used for data simulation.

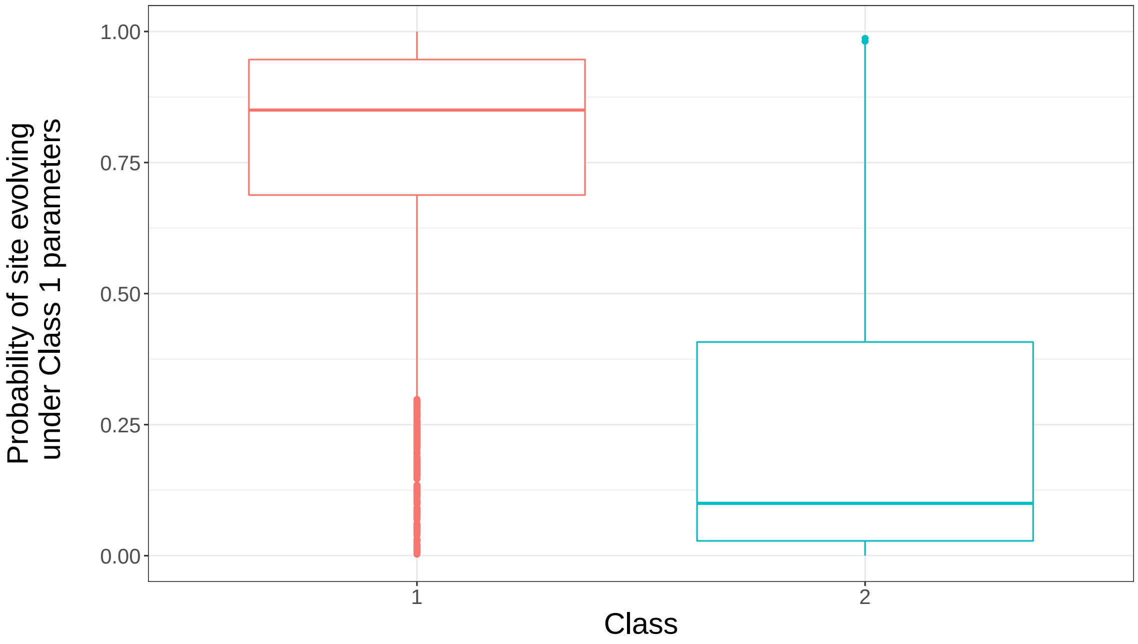


Figure S13: Soft classification of sites to classes - the probability of a site belonging to Class 1 is shown on the y-axis, the two Classes are shown on the x-axis. The boxplots show that in general, sites generated under Class 1 parameters tend to have a higher probability of belonging to Class 1 than sites generated under Class 2.

| RF dist | Number of classes 1 | 2 | 3 | 4 |
|---|---|---|---|---|
| 0 | 116 | 225 | 216 | 208 |
| 2 | 95 | 65 | 68 | 74 |
| 4 | 57 | 9 | 15 | 18 |
| 6 | 20 | 1 | 1 | 0 |
| 8 | 4 | 0 | 0 | 2 |
| $\geq 10$ | 8 | 0 | 0 | 2 |
| Total | 300 | 300 | 300 | 300 |

Table S1: 32-taxon simulations, effect of number of classes inferred on topological accuracy, for the 300 2-class simulated alignments. Columns show the distribution of RF distances, depending on how many classes are inferred. Topological accuracy is greatest when the number of classes inferred matches the number used to simulated the alignments. It is notable that underfitting seems to dramatically decrease topological accuracy, whereas overfitting seems to have a negligible effect in comparison.

| RF dist | Number of classes 1 | 2 | 3 | 4 | 5 |
|---|---|---|---|---|---|
| 0 | 128 | 172 | 201 | 201 | 191 |
| 2 | 101 | 100 | 80 | 78 | 87 |
| 4 | 51 | 24 | 18 | 21 | 20 |
| 6 | 15 | 4 | 1 | 0 | 2 |
| 8 | 3 | 0 | 0 | 0 | 0 |
| 10 | 2 | 0 | 0 | 0 | 0 |
| Total | 300 | 300 | 300 | 300 | 300 |

Table S2: 32-taxon simulations, effect of number of classes inferred on topological accuracy, for the 300 3-class simulated alignments. Columns show the distribution of RF distances, depending on how many classes are inferred. Topological accuracy is greatest when the number of classes inferred matches the number used to simulated the alignments. It is notable that underfitting seems to dramatically decrease topological accuracy, whereas overfitting seems to have a negligible effect in comparison.

| RF dist | Number of classes 1 | 2 | 3 | 4 | 5 | 6 |
|---|---|---|---|---|---|---|
| 0 | 151 | 191 | 203 | 220 | 210 | 205 |
| 2 | 106 | 83 | 82 | 69 | 77 | 80 |
| 4 | 30 | 22 | 13 | 9 | 11 | 12 |
| 6 | 10 | 4 | 1 | 0 | 1 | 2 |
| 8 | 2 | 0 | 1 | 2 | 1 | 1 |
| 10 | 1 | 0 | 0 | 0 | 0 | 0 |
| Total | 300 | 300 | 300 | 300 | 300 | 300 |

Table S3: 32-taxon simulations, effect of number of classes inferred on topological accuracy, for the 300 4-class simulated alignments. Columns show the distribution of RF distances, depending on how many classes are inferred. Topological accuracy is greatest when the number of classes inferred matches the number used to simulated the alignments. It is notable that underfitting seems to dramatically decrease topological accuracy, whereas overfitting seems to have a negligible effect in comparison.

| Model | AIC | BIC |
|---|---|---|
| Discrete Γ | 2,109,008 | 2,109,394 |
| PDF rate | 2,107,607 | 2,108,044 |
| Unlinked GHOST | 2,105,612 | 2,106,900 |

Table S4: Turtle alignment - The AIC and BIC scores returned by the three models tested.

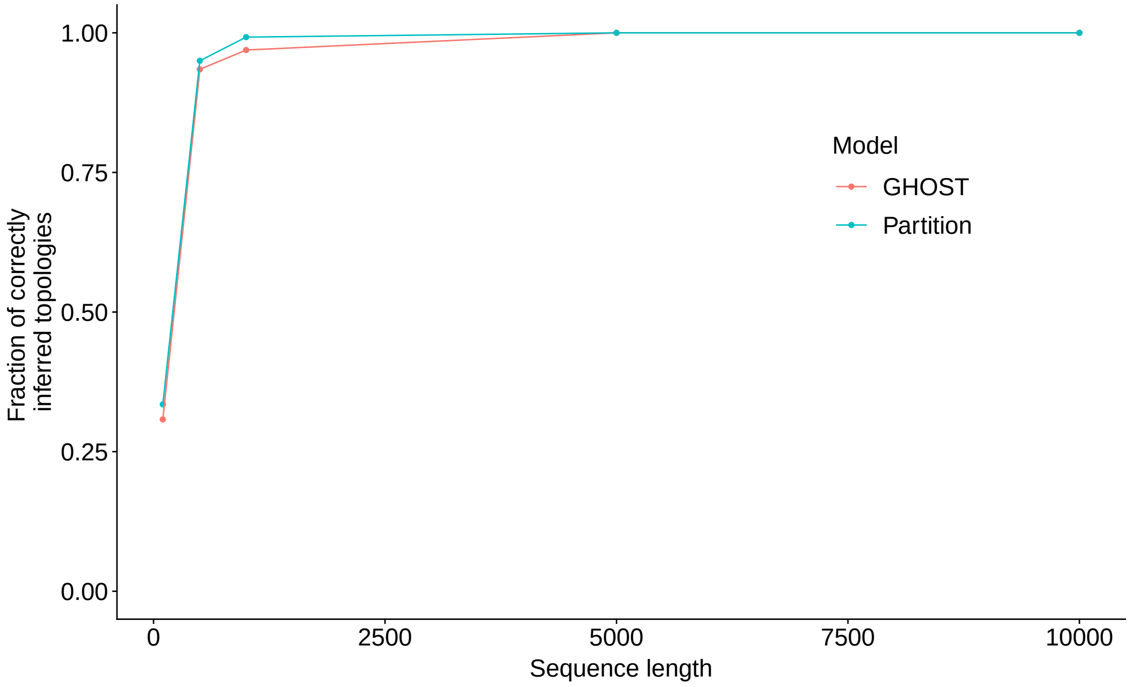


Figure S14: 12-taxon simulations - Proportion of simulated alignments from which the true tree was inferred vs sequence length.

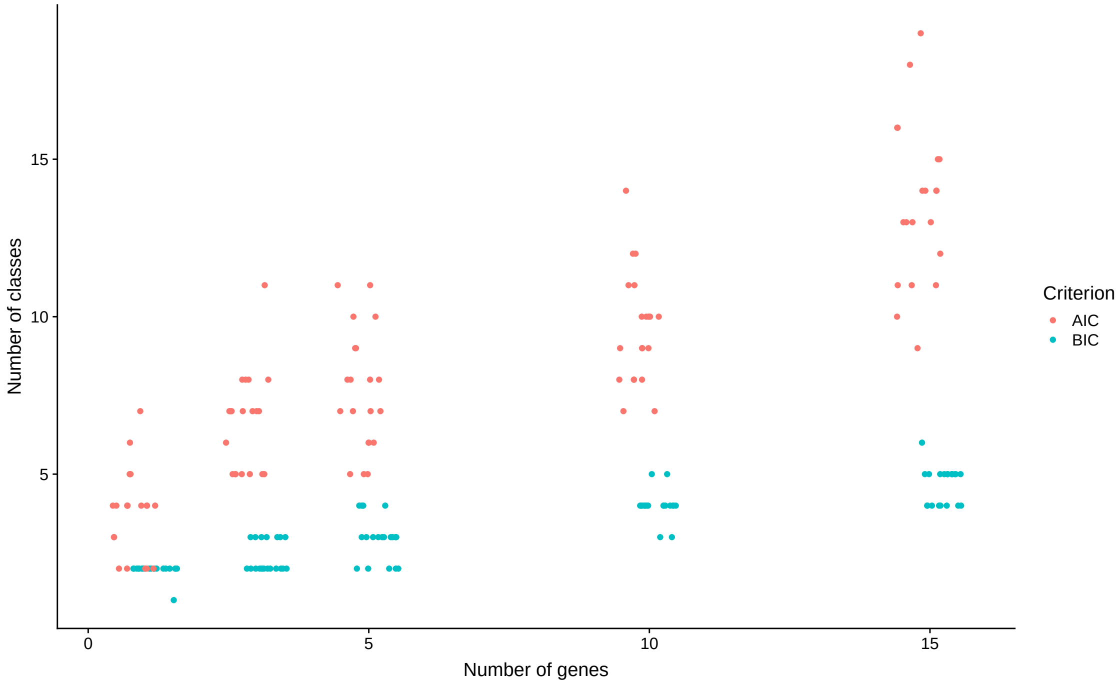


Figure S15: Plastome alignments - number of genes in the alignment vs number of classes preferred by AIC and BIC. Datapoints jittered horizontally for clarity.

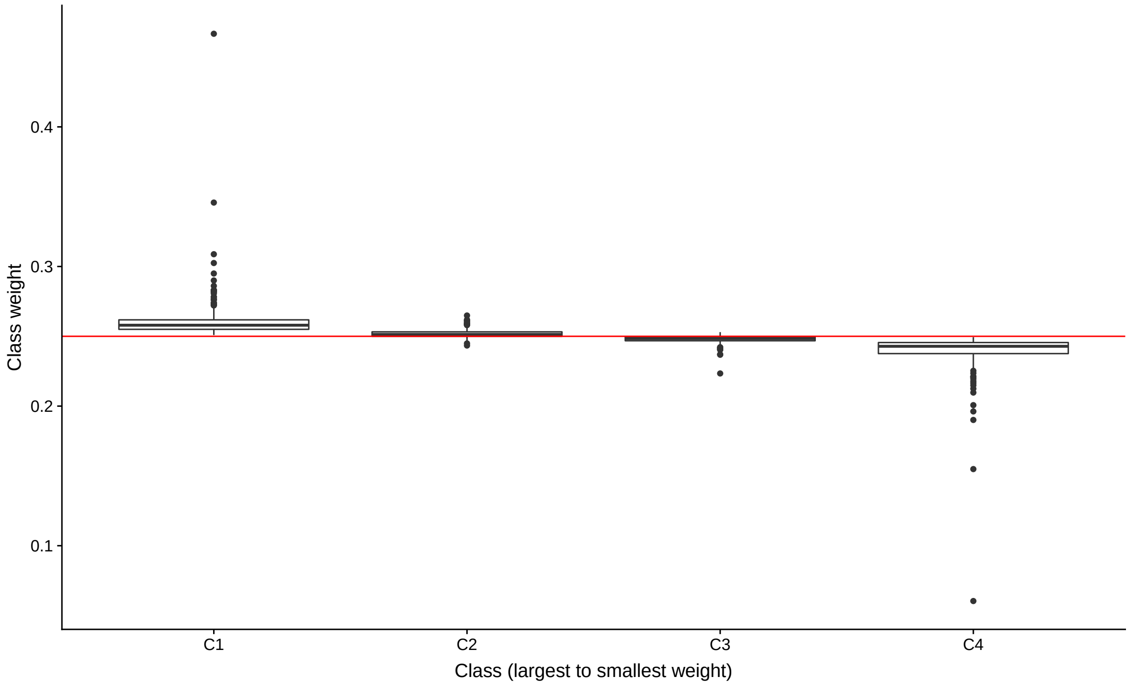


Figure S16: 32-taxon, 4-Class simulations - Weight of inferred classes when the true number of classes (4) is used. The classes are ordered from largest inferred weight (C1) to smallest inferred weight (C4). The red line indicates the true weight of the classes used to simulate the alignments.

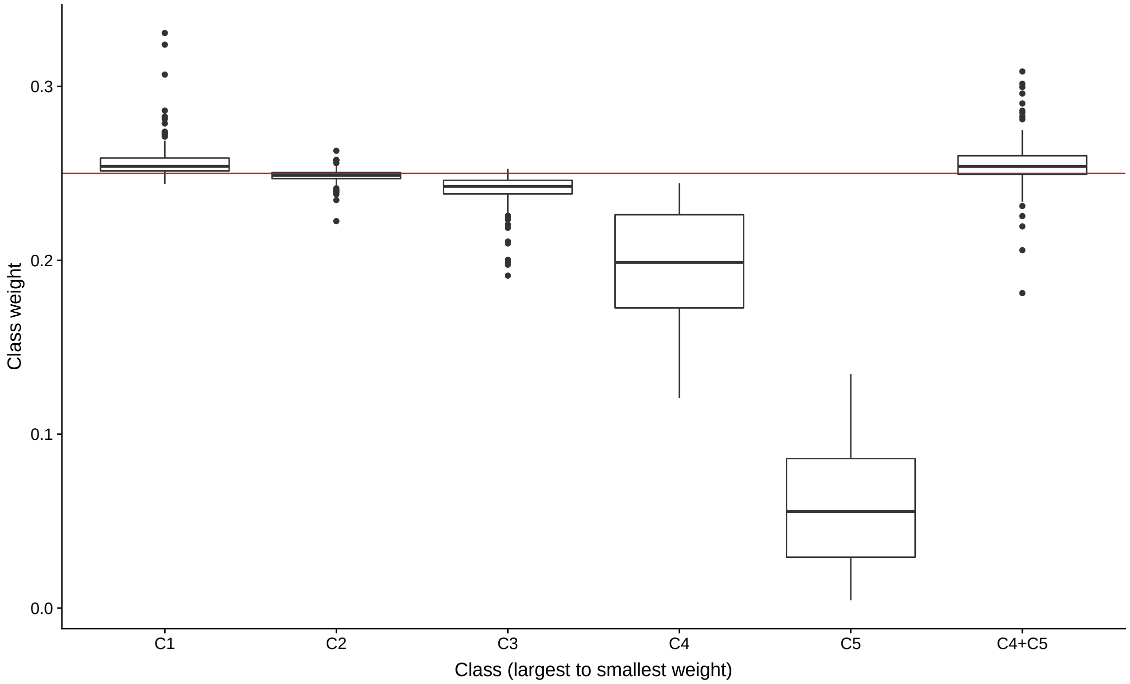


Figure S17: 32-taxon, 4-Class simulations - Weight of inferred classes when overfit by 1 class. The classes are ordered from largest inferred weight (C1) to smallest inferred weight (C5). The red line indicates the true weight of the classes used to simulate the alignments. The final box on the right is the sum of the smallest (by weight) and second smallest classes inferred for each dataset.

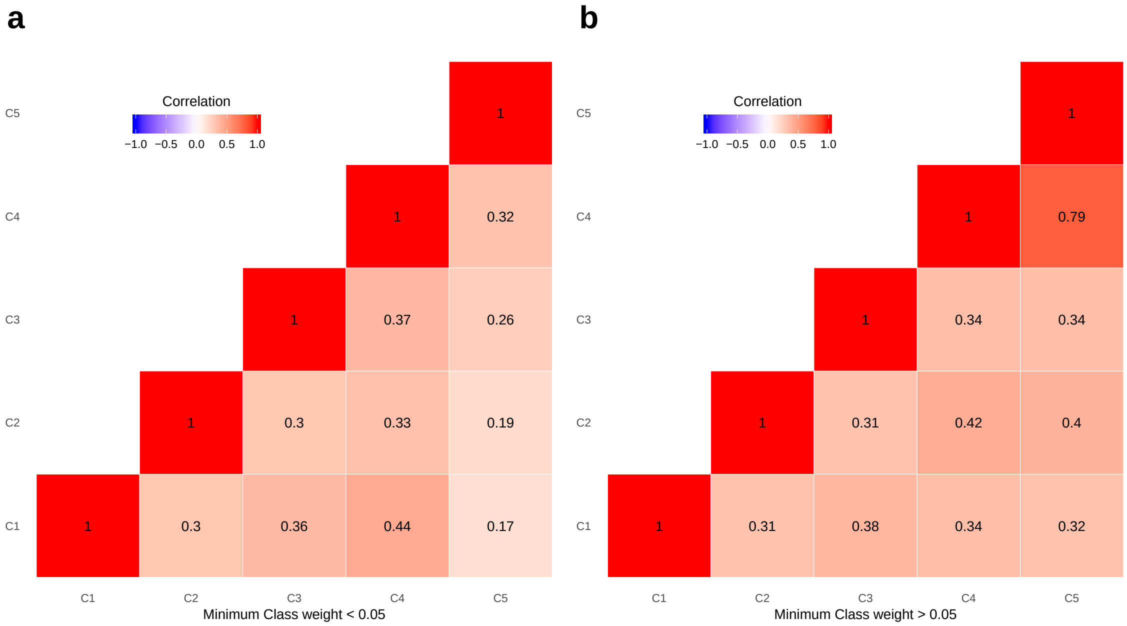


Figure S18: 32-taxon, 4-Class simulations - Correlation of branch lengths between inferred classes when overfit by 1 class. The classes are ordered from largest inferred weight (C1) to smallest inferred weight (C5). (a) The correlation matrix only for those alignments in which the weight of C5 was less than 0.05 (138/300 alignments). (b) The correlation matrix only for those alignments in which the weight of C5 was greater than 0.05 (162/300 alignments).

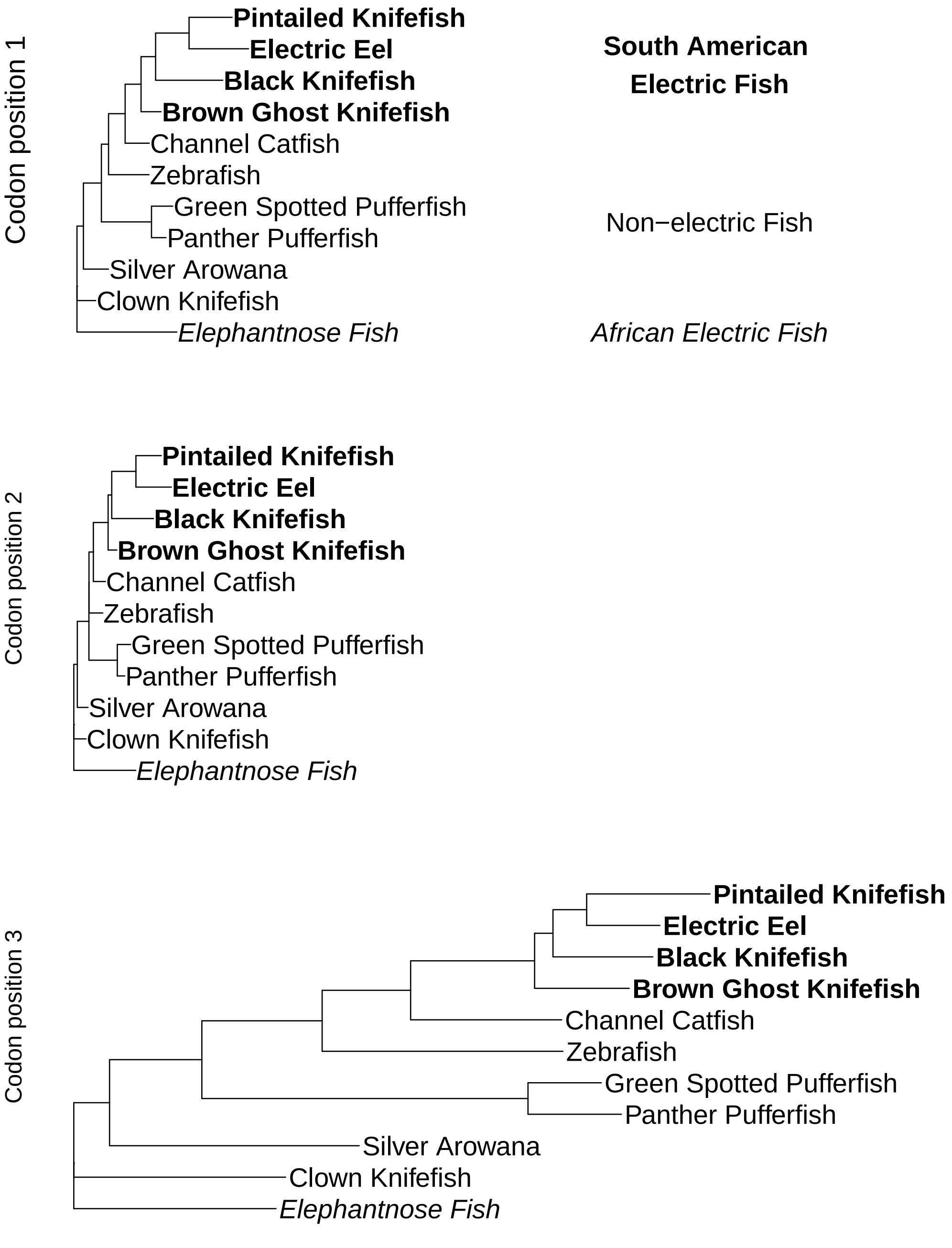


Figure S19: The three trees inferred under the branch-unlinked partition model for the electric fish dataset, with the alignment partitioned based on codon position (CP). The CP1 and CP2 partitions used a GTR+FO+G model, while the CP3 partition used a GTR+FO+I+G model.

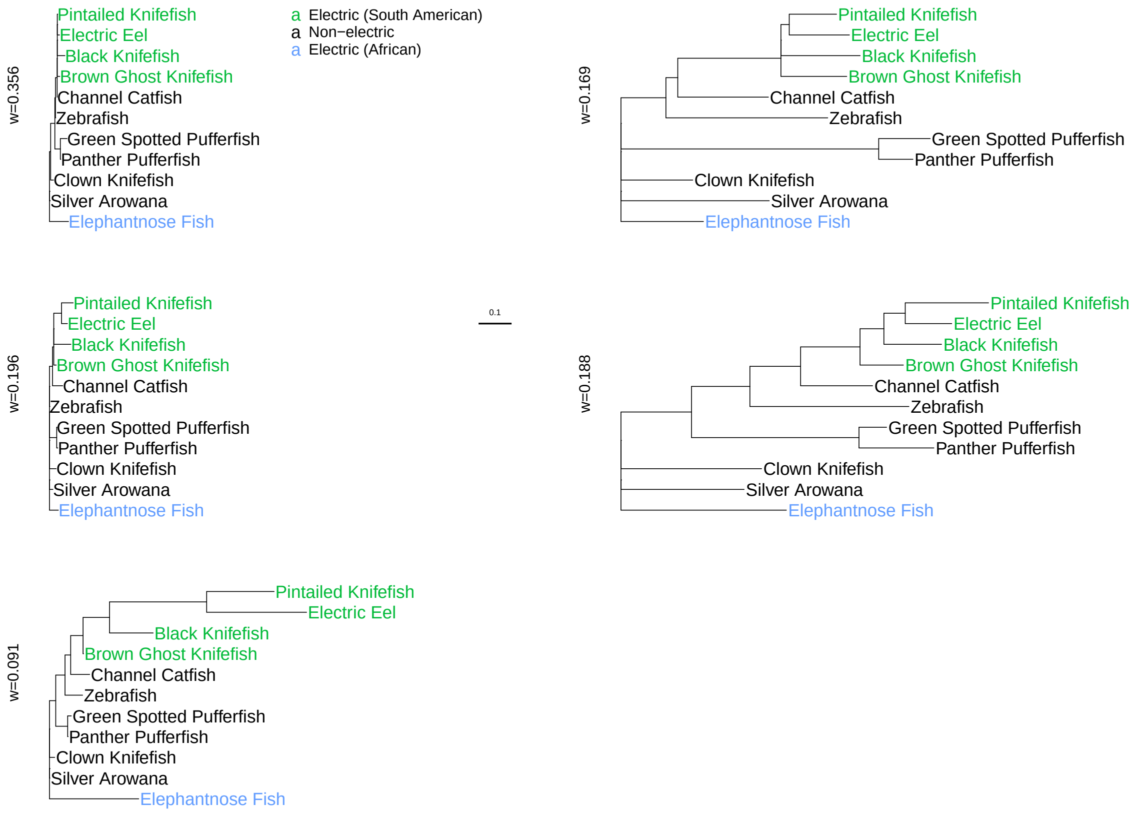


Figure S20: The five trees inferred under the General Time Reversible, five class mixture model (GTR+FO*H5) for the electric fish data.

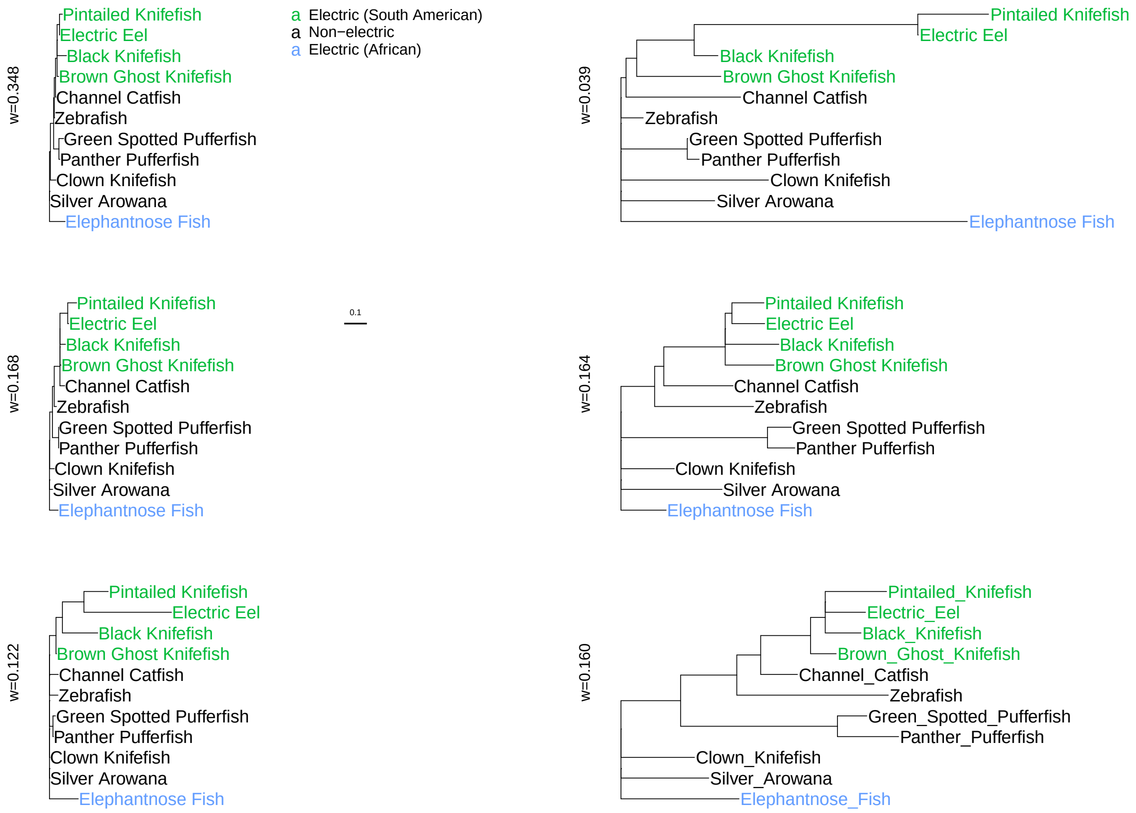


Figure S21: The six trees inferred under the General Time Reversible, six class mixture model (GTR+FO*H6) for the electric fish data.
